# Local fractal dimension as an indicator of functional reorganization in molecular interaction networks

**DOI:** 10.64898/2026.10.27.762149

**Authors:** Frank Hause, Alice Wedler-Geyer, Marcell Lederer, Tommy Fuchs, Christian H. Ihling, Dirk Tänzler, René Keil, Oleksandr Sorokin, Sebastian Wussow, Andrea Sinz, Stefan Hüttelmaier

## Abstract

Changes in protein abundance incompletely describe how molecular perturbations reshape cellular organization. Here we introduce DIFFRACT, a framework that integrates quantitative proteomics with protein-protein interaction networks to identify reorganization through changes in local fractal dimension. Molecular associations define effective distances between connected proteins, allowing the geometry of their surrounding network neighborhoods to be compared across conditions. Analyses across four organisms (*E. coli, A. thaliana*, *C. elegans*, and *H. sapiens*) and three interaction resources (STRING, BioGRID, and IntAct) establish heterogeneous local scaling, while controlled perturbations demonstrate sensitivity to regional changes and invariance to uniform distance rescaling. Applied to RAVER1-deficient A549 cells, DIFFRACT identifies components associated with established RNA-regulatory functions and candidate roles in tRNA biology, mitochondrial organization and ciliogenesis. Selected RNA-regulatory and ciliary themes are not significantly enriched by abundance-ranked gene set analysis, supporting complementary information from network geometry. Selected functional annotations also recur in HEK293T cells. Guided by the ciliary predictions, we show that RAVER1 depletion increases the proportion of ciliated A549 cells 2.57-fold and increases cilium length. DIFFRACT thus connects changes in molecular network geometry to experimentally testable biological responses, extending the interpretation of proteomic perturbations beyond differential abundance.

## Introduction

Molecular profiling provides increasingly comprehensive measurements of cellular composition, but abundance changes alone do not explain how perturbations reorganize biological processes. Systems biology addresses this gap by considering the interactions through which molecular components contribute to cellular function (Kitano, 2002; Barabási & Oltvai, 2004; Joyce & Palsson, 2006). Integrating molecular profiles with interaction networks can reveal coordinated responses that are incompletely captured by testing individual genes or proteins and annotating the resulting lists (Mitra et al., 2013).

Interaction resources such as BioGRID (Stark et al., 2006), IntAct (del Toro et al., 2022) and STRING (von Mering et al., 2003; Szklarczyk et al., 2025) provide scaffolds onto which molecular measurements can be mapped. On a defined scaffold, molecular associations can be represented as effective distances: stronger coordination corresponds to shorter distances, whereas weaker coordination corresponds to longer distances. Condition-dependent weighting therefore changes the network metric and the accessibility of surrounding proteins. Quantifying this organization requires a readout that integrates relationships across a protein’s neighborhood.

Local fractal dimension (LFD) describes how the cumulative neighborhood around a node grows with network distance (Pu et al., 2014). It builds on the observation that complex networks can exhibit self-similar organization over finite distance ranges and heterogeneous scaling across regions (Song et al., 2005; Furuya & Yakubo, 2011). When topology is held fixed, changes in LFD describe geometric consequences of altered edge distances, including effects extending beyond immediate neighbors.

Here, we introduce DIFFRACT (Differential Fractal Analysis of Condition-Specific Topologies), a framework integrating replicated molecular-abundance profiles with interaction scaffolds. DIFFRACT derives effective edge distances from molecular associations and separates weighted LFD into a structural baseline and a weighting-dependent contribution. Differences in the latter prioritize proteins with unusual changes in neighborhood geometry; connected components of these proteins provide candidates for functional annotation.

We examine network scaling across four organisms (*E. coli, A. thaliana*, *C. elegans*, and *H. sapiens*) and three interaction resources (STRING, BioGRID, and IntAct), test LFD responses to controlled edge rescaling, and compare proteome-derived weighting strategies against expression-permuted controls. Applying DIFFRACT to RAVER1-deficient A549 cells recovers established RNA-regulatory functions and identifies candidate processes not highlighted by abundance-based enrichment, i.e. overrepresentation analysis (ORA) (Draghici et al., 2003) or gene set enrichment analysis (GSEA) (Subramanian et al., 2005). Selected functional annotations recur in HEK293T cells, and targeted cell culture experiments reveal increased ciliation and cilium length following RAVER1 depletion. These analyses position differential LFD as a complementary means of interpreting molecular perturbations and generating experimentally testable hypotheses.

## Results

### Molecular interaction networks exhibit conserved multifractal organization

The use of LFD to quantify molecular network reorganization requires interaction networks to exhibit scaling over finite topological distances and sufficient local heterogeneity to support node-resolved analysis. We therefore first examined the structural organization of physical protein-protein interaction (PPI) networks from *Escherichia coli*, *Arabidopsis thaliana*, *Caenorhabditis elegans* and *Homo sapiens*. To assess whether the observed properties were robust to differences in network coverage and curation, we independently analyzed networks obtained from BioGRID (Stark et al., 2006), IntAct (del Toro et al., 2022) and STRING (von Mering et al., 2003; Szklarczyk et al., 2025).

Box-covering analysis revealed an approximately linear decrease in the normalized number of boxes, N_B_(l_B_)/N, with increasing box size, l_B_, on logarithmic axes over defined scaling intervals (Fig. 1a). The corresponding box-counting dimensions varied between organisms and interaction resources. For *E. coli*, d_B_ was 5.22, 5.30 and 1.76 for BioGRID, IntAct and STRING, respectively. The corresponding estimates were 6.80, 6.85 and 1.70 for *A. thaliana*; 5.16, 6.95 and 1.52 for *C. elegans*; 7.49, 6.52, and 5.37 for the human PPI networks. At larger box sizes, the curves approached a plateau as the finite networks became coverable by only a small number of boxes. These results identify finite scaling regimes across organisms while also demonstrating substantial resource-dependent variation in their estimated box-counting dimensions.

The corresponding multifractal spectra were broad and non-degenerate rather than collapsing onto a single scaling exponent (Fig. 1b). This indicated heterogeneous local scaling within each network, with different regions contributing distinct Lipschitz-Hölder exponents. Spectrum position and width differed between interaction resources, consistent with differences in network coverage and curation, but multifractal organization was retained across all four organisms. Together, these findings show that molecular interaction networks possess reproducible global scaling alongside heterogeneous local geometry, providing the structural basis for resolving condition-dependent changes through node-wise fractal dimensions.

**Figure 1.**
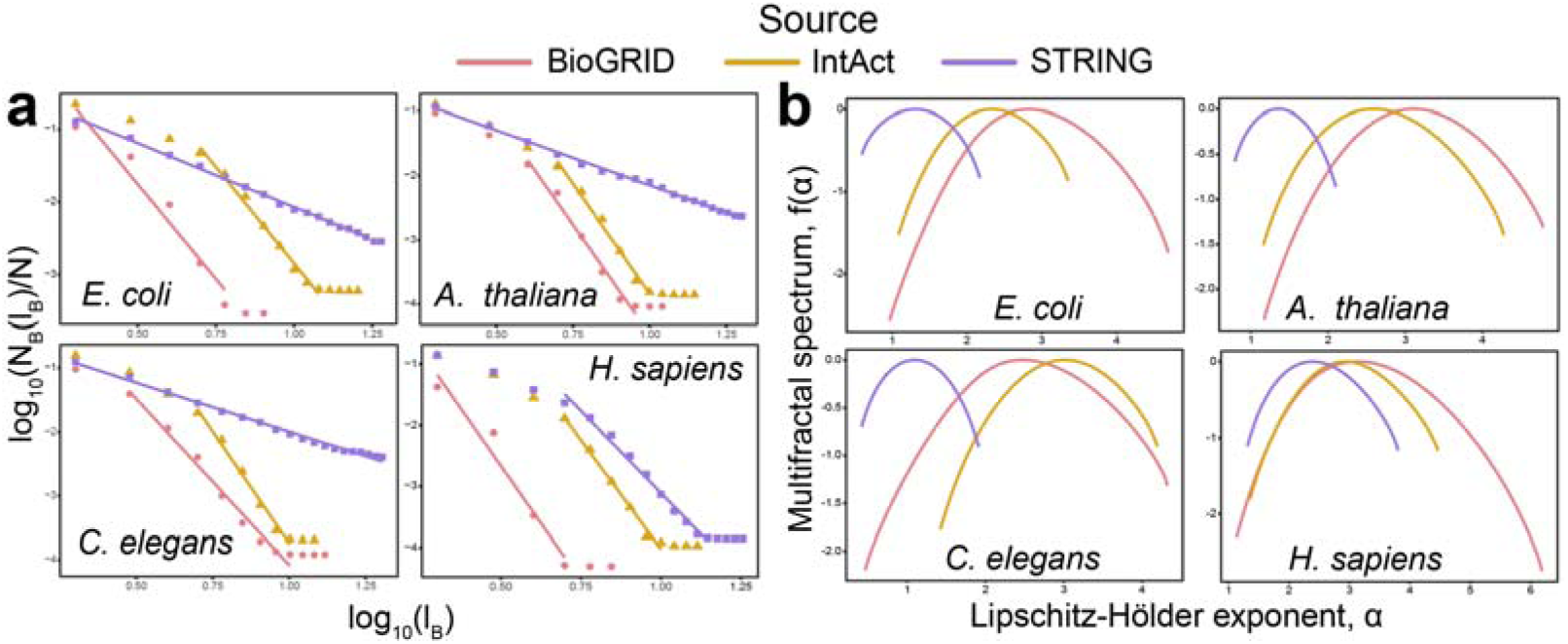
Molecular interaction networks exhibit fractal scaling and heterogeneous local geometry across organisms and interaction resources. **a)** Box-covering analysis of molecular interaction networks from *E. coli*, *A. thaliana*, *C. elegans* and *H. sapiens* obtained from BioGRID (pink circles), IntAct (ochre triangles) and STRING (purple squares). The normalized number of boxes required to cover each network, N_B_(l_B_)/N, is plotted against box size l_B_ on logarithmic axes. Solid lines denote linear fits within the selected scaling intervals. The approximately linear relationships are consistent with fractal scaling over finite ranges; plateaus at larger box sizes reflect finite-network saturation. b) Multifractal spectra, f(α), for the corresponding networks, where α denotes the Lipschitz-Hölder exponent. The broad spectra demonstrate heterogeneous local scaling, whereas differences in spectrum position and width indicate variation in scaling organization among interaction resources and organisms. Colors denote interaction resources as in a).

### Local fractal dimensions selectively resolve regional changes in network geometry

Having established that molecular interaction networks exhibit heterogeneous fractal organization, we next tested whether LFD can distinguish local changes in network geometry from differences arising from network structure or arbitrary distance scaling. Across the 12 organism-resource combinations, empirical networks exhibited distinct median LFD values that were not reproduced by an ensemble of degree-preserving null networks, confirming that LFD captures organization beyond the degree distribution alone (Fig. 2a).

A valid measure of local geometry should remain unchanged when all edge distances are multiplied by the same constant, as such global rescaling changes only the unit of distance and not the relative organization of the network. Accordingly, multiplying all edge distances by factors ranging from 0.1 to 10 produced maximum absolute LFD differences of only approximately 10^-15^ to 10^-14^, consistent with numerical precision (Fig. 2b). By contrast, rescaling only the edges within defined connected regions produced LFD changes approximately 14 orders of magnitude larger than those observed after global rescaling (Fig. 2c). Thus, LFD was invariant to a change in global distance scale but responsive to regional alterations in the network metric.

To characterize this response systematically, we selected three connected 100-node regions within each network and multiplied their internal edge distances by factors ranging from 0.25 to 4 while retaining the nodes, edges and all distances outside the respective region. Rescaling factors < 1 modelled local contraction, whereas factors > 1 modelled local relaxation. Regional rescaling produced factor-dependent changes in LFD across organisms and interaction resources (Fig. 2d). The largest signed responses generally occurred within the perturbed regions, with smaller responses among adjacent and more distant nodes. In a representative region of the human BioGRID network, halving internal edge distances produced a predominantly negative LFD response within the perturbed region, whereas doubling them produced the opposite response (mean ΔLFD = -1.69 and 2.60, respectively; Fig. 2e).

Although strongest within and immediately surrounding the perturbed regions, changes in LFD were not restricted to rescaled nodes. Stratification by shortest-path distance showed that the response propagated through the wider network, with its magnitude and spatial extent depending on the organism, interaction resource and direction of rescaling (Fig. 2f). Together, these controlled perturbations establish that LFD measures relative, spatially distributed changes in network geometry rather than absolute edge scale. This scale invariance and regional sensitivity provide the metric basis for using differential LFD to quantify condition-specific network reorganization.

**Figure 2.**
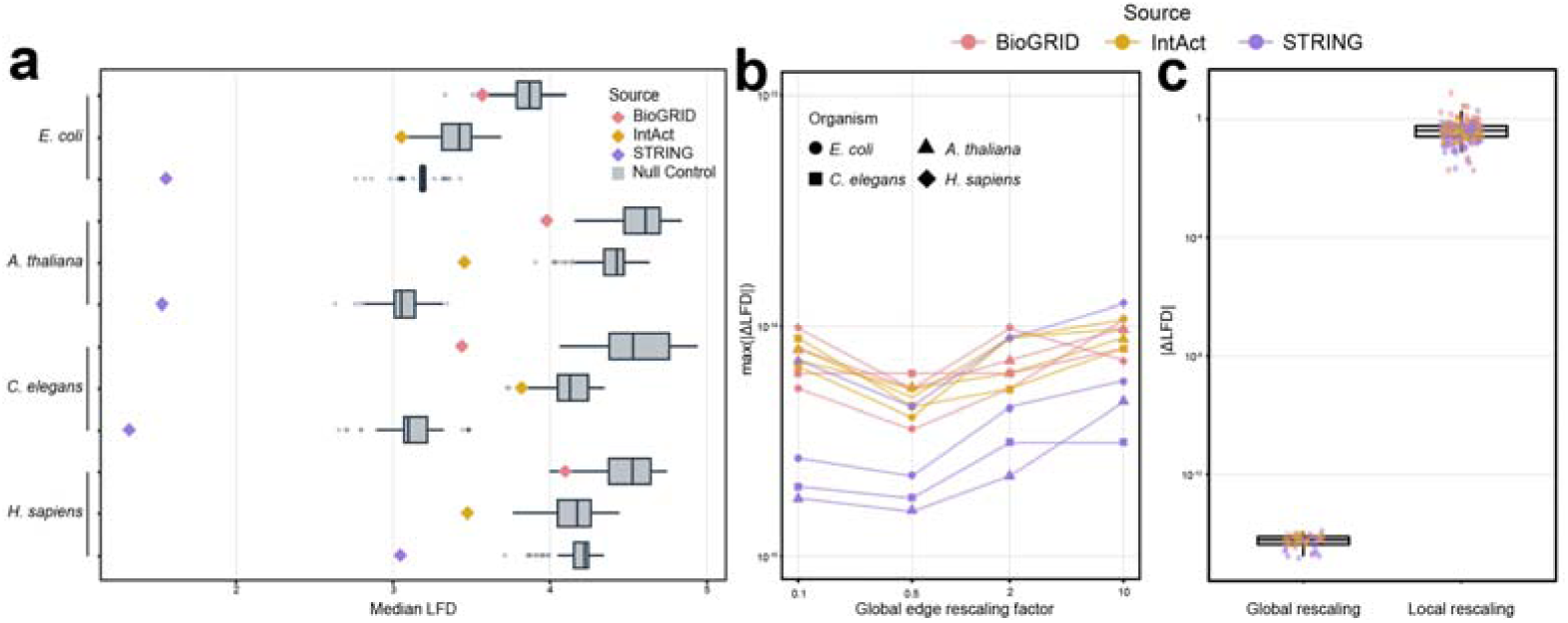
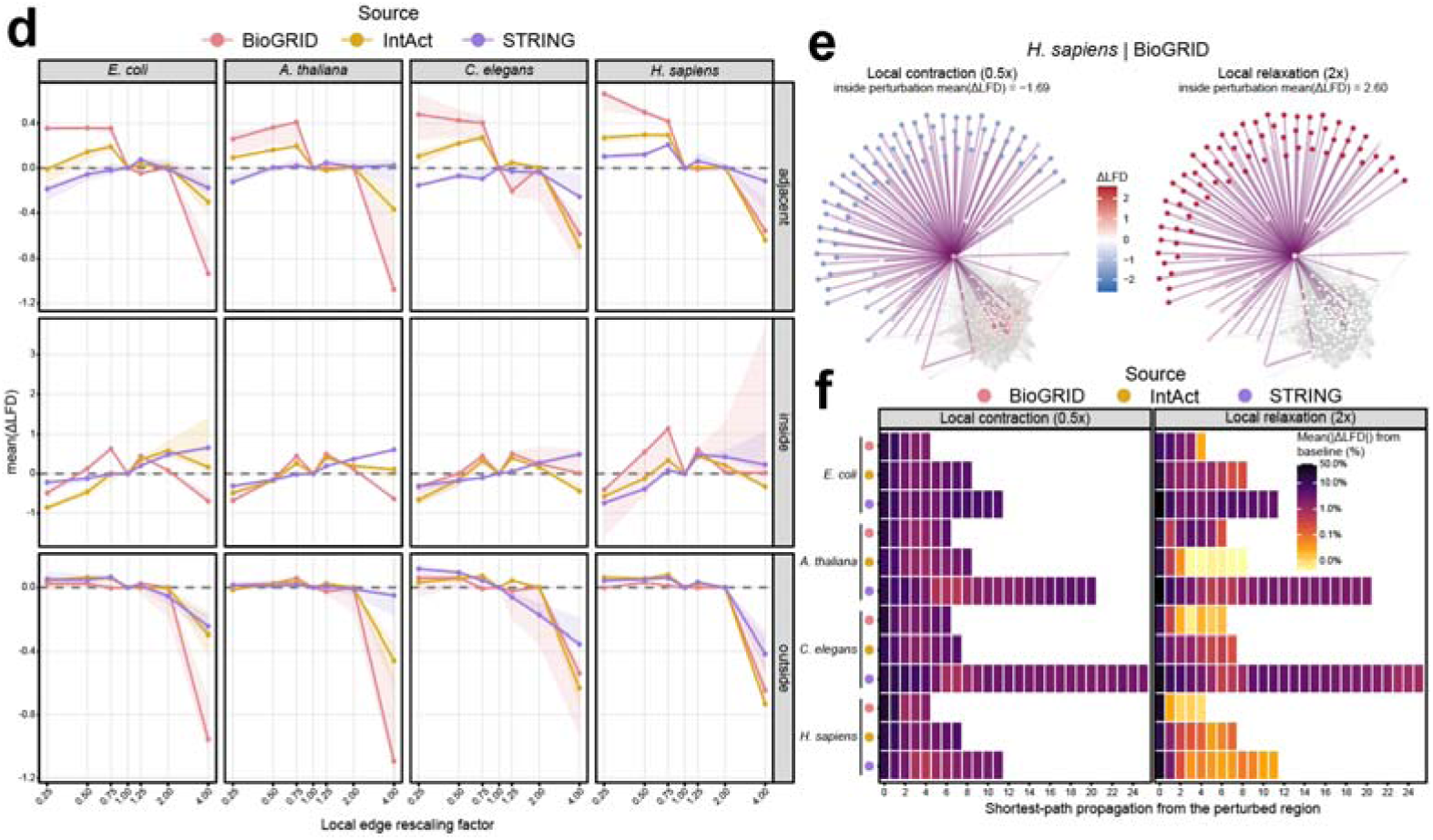
Local fractal dimensions are invariant to global distance scaling but respond to regional network contraction and relaxation. **a)** Median structural LFD across molecular interaction networks from *E. coli*, *A. thaliana*, *C. elegans* and *H. sapiens*. Colored diamonds show empirical networks from BioGRID, IntAct and STRING. Grey box plots show degree-preserving null ensembles (n = 1,000 randomized networks per empirical network); center lines denote medians, boxes the interquartile range and whiskers values within 1.5 times the interquartile range. **b)** Maximum absolute change in node-wise LFD after multiplying all edge distances by the indicated global scaling factor. Colors denote interaction resources and symbols denote organisms. Differences remained at numerical precision across scaling factors. **c)** Absolute LFD responses following global or local edge-distance rescaling. Local responses were calculated for nodes within the controlled regions across non-baseline perturbation factors. Points denote individual organism-, resource- and perturbation-specific measurements; colors denote interaction resources. **d)** Median signed change in LFD after rescaling the internal edge distances of connected 100-node regions by the indicated factor. Factors < 1 represent local contraction, 1 is the unperturbed baseline and factors > 1 represent local relaxation. Columns denote organisms and rows classify nodes as adjacent to, inside or outside the perturbed region. Lines show medians and shaded areas summarize variation across three independently selected regions; colors denote interaction resources. **e)** Node-wise ΔLFD in a representative 100-node region of the human BioGRID network after local contraction (0.5-fold internal edge distance, left) or relaxation (2-fold internal edge distance, right). Pink edges indicate the rescaled internal edges and grey edges the surrounding network. Nodes are coloured by ΔLFD. **f)** Spatial propagation of the LFD response following contraction or relaxation of a fixed 100-node region. Color represents the mean absolute percentage change in LFD relative to the unperturbed baseline, stratified by shortest-path distance from the perturbed region. Rows denote organism-interaction-resource combinations.

### Mutual information-weighting captures non-random proteomic organization in interaction networks

The cross-species analysis established that fractal organization is a general property of molecular interaction networks. Condition-specific edge weighting, however, requires replicated molecular abundance measurements matched to the organism and biological system under study. As experimental application, proteome reorganization was examined in human lung cancer A549 cells restricting the development and validation of the weighting strategy to human interaction networks. Retaining BioGRID, IntAct and STRING as independent interaction scaffolds allowed us to assess whether the resulting network geometry was robust to differences in human interactome coverage and curation.

We next sought to derive biologically informed edge distances from molecular abundance data. Quantitative proteomic profiles from three wild-type (WT) replicates of the human lung adenocarcinoma cell line A549 were integrated with fixed BioGRID, IntAct and STRING interaction scaffolds to construct condition-specific weighted networks. Edge associations were quantified using mutual information (MI), absolute Pearson correlation or distance correlation (dCor). We additionally included a variance/missingness weighting, termed VarMiss, in which edge distances were determined by marginal protein variance and pairwise data completeness without edge-specific dependence information. As a null model, protein abundance values were independently permuted across samples while preserving the observed zero and missing-value masks.

Expression-derived edge-weight distributions differed from their permutation controls for all three dependence measures, whereas VarMiss retained only the contribution of marginal variance and data completeness (Fig. 3a). We then decomposed node-wise LFD into a structural component determined by the unweighted interaction scaffold and a dynamic component introduced by abundance-dependent edge weighting. Across interaction resources, physiological expression weighting shifted dynamic LFD distributions away from their permutation controls and produced spatially structured changes across the networks (Fig. 3b).

Total LFD was consistently higher under physiological expression weighting than under either expression-permuted weighting or the corresponding unweighted topology for all combinations of interaction resource and weighting method (Fig. 3c). For MI weighting, the median node-wise difference between physiological and expression-permuted total LFD was 1.086 in BioGRID, 0.381 in IntAct and 0.435 in STRING. Relative to the unweighted structural networks, the corresponding differences were 1.238, 0.572 and 0.550. Pearson correlation and dCor produced changes of similar direction, whereas the VarMiss control generally showed smaller expression-dependent effects, particularly in BioGRID and IntAct.

The different weighting strategies produced broadly comparable node-wise LFD fit quality, with median adjusted (R^2^) values of 0.479–0.798 and median root-mean-square errors (RMSE) of 0.560–0.722 (Fig. 3d). Spatial continuity depended more strongly on the underlying interaction resource than on the association measure: adjacent-node dynamic LFD correlations were slightly negative in BioGRID, intermediate in IntAct and highest in STRING. Conversely, mean absolute differences between adjacent nodes decreased from BioGRID to STRING. These observations indicate that the structural interaction scaffold constrains the spatial organization of the resulting dynamic LFD field, while the expression-derived weights determine its displacement from the structural baseline.

To compare the ability of the weighting methods to separate physiological organization from randomized expression, we combined the directionally harmonized permutation contrasts into a standardized null-discrimination score. MI achieved the highest aggregate score for each interaction resource (BioGRID, 0.60; IntAct, 0.43; STRING, 0.55), whereas Pearson and dCor showed weaker and less consistent separation and VarMiss yielded negative scores in all three networks (Fig. 3e). We therefore selected MI as the primary metric for deriving expression-informed edge weights in subsequent analyses. This choice was based on its reproducible discrimination of physiological from permuted network geometry across different interaction resources, rather than on differences in LFD fit quality alone.

**Figure 3.**
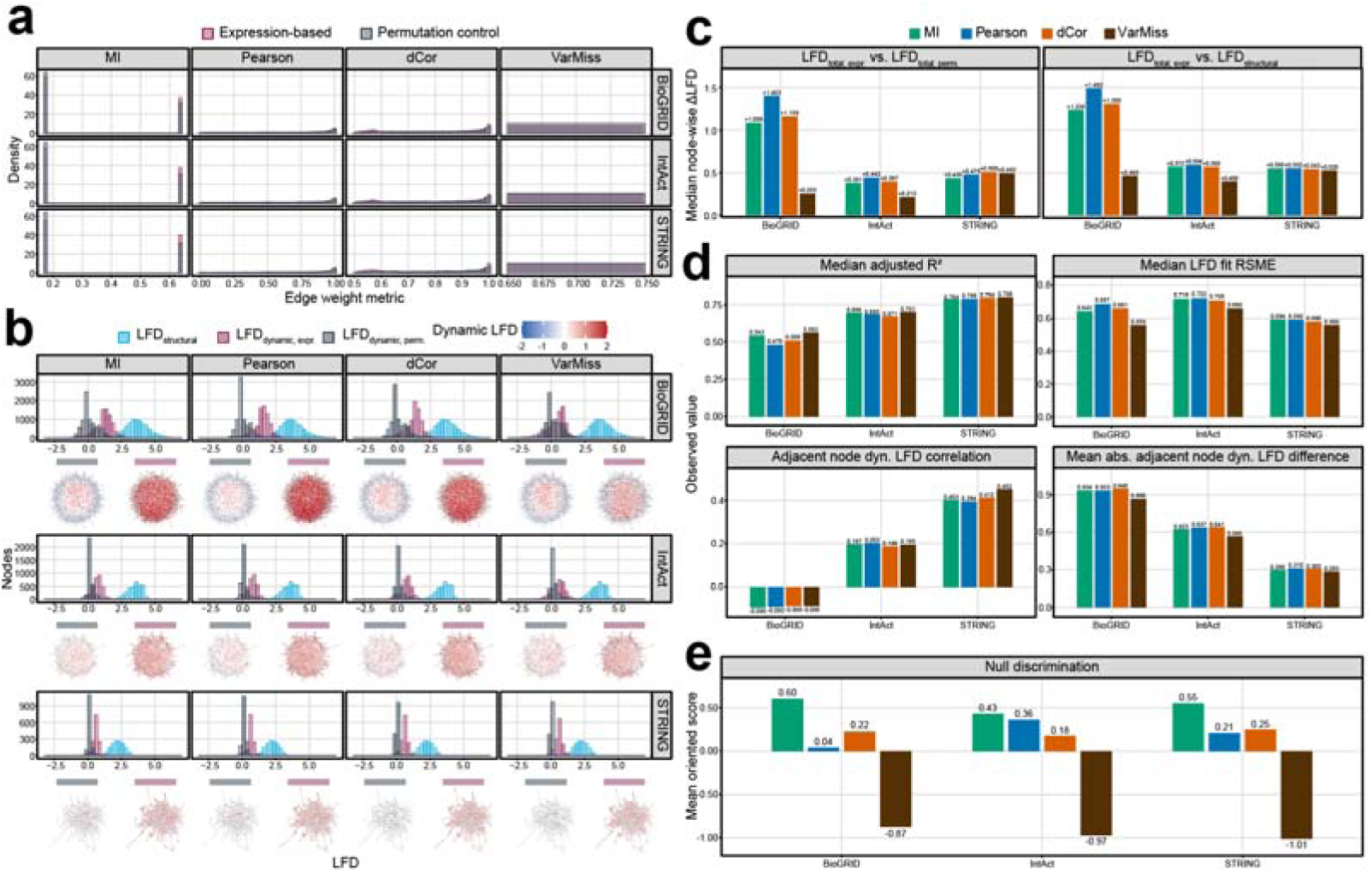
Mutual information-weighting transforms proteomic covariation into non-random local network geometry. **a)** Distributions of edge-weight metrics derived from the measured proteomic profiles of WT A549 cells and their expression-permuted controls across BioGRID, IntAct and STRING interaction networks. Edge associations were quantified using mutual information (MI), absolute Pearson correlation or distance correlation (dCor). VarMiss denotes a variance/missingness weighting, in which edge weights depend on marginal protein variance and pairwise data completeness but not on edge-specific molecular dependence. **b)** Node-wise LFD decomposition and corresponding network projections for each interaction resource and weighting method. Total weighted LFD was decomposed into structural and dynamic components, with dynamic LFD defined as LFD_dynamic_ *=* LFD_weighted_ - LFD_structural_. Distributions show structural LFD, expression-derived dynamic LFD and permutation-control dynamic LFD. Network layouts were held constant within each interaction resource; node colours indicate dynamic LFD. **c)** Median node-wise differences in total LFD between physiological expression-weighted and expression-permuted networks (left) and between physiological expression-weighted and unweighted structural networks (right). Values above bars indicate the median ΔLFD. **d)** Diagnostic properties of expression-weighted LFD estimates. Panels show the median adjusted (R^2^) and root-mean-square error (RMSE) of node-wise LFD fits, the correlation of dynamic LFD between adjacent nodes and the mean absolute dynamic LFD difference between adjacent nodes. Values are shown separately for BioGRID, IntAct and STRING and for each weighting method. **e)** Aggregate null-discrimination scores derived from standardized, directionally harmonized contrasts between physiological expression-weighted and expression-permuted networks. Higher values indicate stronger separation from the permutation control. MI produced the highest score for all three interaction resources. Proteomic weighting used three WT A549 samples; MI was estimated with two equal-frequency bins.

### RAVER1 depletion reorganizes functional network modules beyond differential protein abundance

We applied DIFFRACT to three wild-type (WT) and three RAVER1-knockout (KO) A549 proteomes. Principal component analysis separated the conditions (Fig. 4a), whereas differential abundance analysis identified only 11 proteins meeting both thresholds (adjusted P < 0.05; |log₂(FC)| > 1), with three increased and eight decreased (Fig. 4b). We therefore asked whether network geometry could reveal broader functional responses.

Condition-specific edge distances were calculated independently from WT and KO profiles. Node-wise ΔLFD was defined as the KO–WT difference in dynamic LFD, and proteins with network-relative outlier scores were mapped onto each scaffold. Selected proteins were grouped into connected components: sets in which every protein can be reached from every other through scaffold links involving only selected proteins. Within each scaffold– weighting analysis, these components contain distinct, non-overlapping protein sets, although their biological functions may overlap. Components containing at least five proteins were retained. MI was the primary weighting metric; Pearson correlation, distance correlation and VarMiss provided complementary comparisons (Fig. 4c).

The preceding analyses used physical interaction networks. Because these maps incompletely capture relationships among proteins contributing to shared processes, we additionally examined a high-confidence STRING physical-plus-functional scaffold for functional interpretation. MI weighting yielded a dominant 305-protein component in BioGRID and components of 21 and 12 proteins in IntAct and physical STRING, respectively. On the expanded STRING scaffold, selected outliers formed ten components containing 5–15 proteins, enabling individual annotation.

Component enrichment revealed cell–extracellular matrix interactions, junction organization and cell communication in IntAct, and microtubule-dependent transport, centrosome organization, mitotic progression and nucleotide metabolism in physical STRING (Supplementary Fig. S1). The expanded STRING scaffold distinguished components associated with RNA regulation, tRNA modification, amino-acid metabolism, vesicle trafficking, apoptosis and mitochondrial organization. Its largest MI-derived component included enrichment for miRNA-mediated gene silencing, RISC assembly and pre-miRNA processing, consistent with established RAVER1 functions. These findings illustrate how geometric outlier selection organizes the proteomic response into functionally interpretable candidate modules beyond the limited set of differentially abundant proteins.

**Figure 4.**
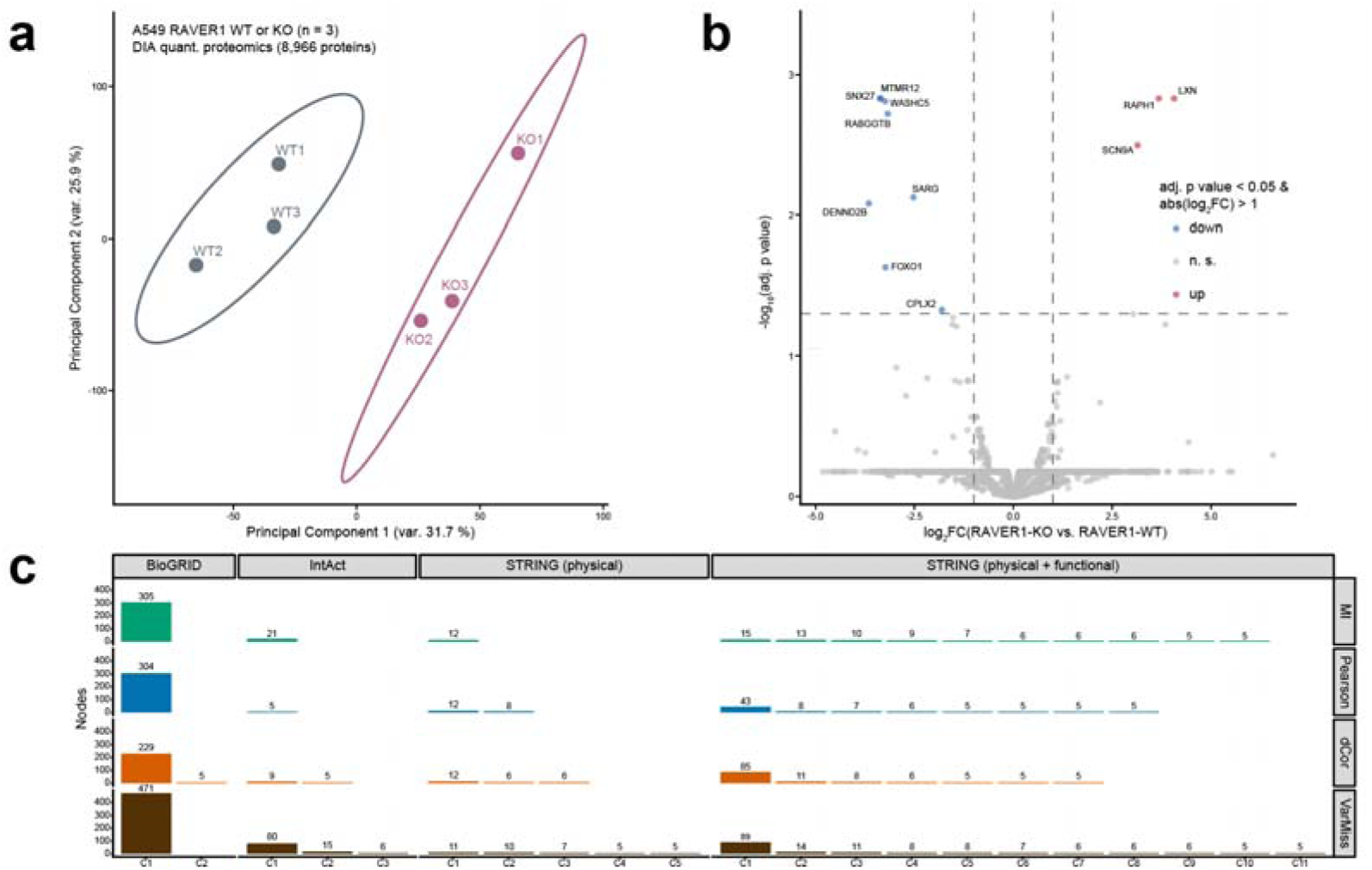
Changes in local fractal geometry reveal functional network reorganization following RAVER1 deletion. **a)** Principal component analysis of quantitative proteomic profiles from WT and RAVER1-KO A549 cells (each n = 3). The analysis included 8,966 quantified proteins. Labels identify individual replicates, and ellipses denote 95% confidence regions. Percentages indicate the variance explained by each principal component. **b)** Volcano plot of differential protein abundance between RAVER1-KO and WT cells. Proteins were classified as significantly increased or decreased at an adj. p-value < 0.05 and an absolute log_2_(FC) > 1. Dashed lines indicate the significance and fold-change thresholds. Selected proteins with the strongest evidence for differential abundance are labelled. RAVER1 was not detected in the KO condition, hence, it is not present in the Volcano plot. **c)** Numbers of proteins in connected components formed by network-relative ΔLFD outliers between RAVER1-KO and WT cells. Each bar represents a group of selected proteins connected directly or indirectly through other selected proteins on the indicated interaction scaffold; only groups containing at least five proteins are shown. Components within each panel do not share proteins, but protein membership can overlap across panels representing different scaffolds or weighting methods. Columns denote BioGRID, IntAct, physical STRING and high-confidence STRING physical-plus-functional scaffolds. Rows denote MI, absolute Pearson correlation, distance correlation (dCor) and variance/missingness weighting (VarMiss). Component labels (C1, C2, etc.) are assigned separately within each panel and do not imply corresponding membership across panels. Numbers above bars indicate protein counts.

### Changes in fractal network geometry reveal established and candidate roles of RAVER1

To interpret the response to RAVER1 deletion, we analysed the proteins in each of the ten MI-derived components from the physical-plus-functional STRING network separately for overrepresentation of Gene Ontology (Ashburner et al., 2000; The Gene Ontology Consortium, 2023) and Reactome biological processes and Reactome (Milacic et al., 2024) pathways. This analysis identified 351 significant component–term associations, indicating processes or pathways represented more frequently among a component’s proteins than expected from the analysis background. Because related annotations often describe the same proteins, we grouped annotations supported by overlapping protein sets within each component into 38 functional themes. These themes summarize the enrichment results without introducing additional statistical tests.

We further assessed the sensitivity of each theme by repeating the enrichment analysis after removing individual component proteins while preserving the annotation universe and multiple-testing family. Sixteen themes remained significant after every single-protein removal, identifying functional signals that were not attributable to an individual protein.

Several components recovered functions previously linked to RAVER1 (Fig. 5a). C2 was dominated by a robust RNA-splicing theme comprising 11 spliceosomal or spliceosome-associated proteins, including SF1, CDC5L, CWC22, ISY1, SUGP1 and PSME3IP1. This organization was consistent with the established function of RAVER1 as a PTBP1-associated splicing co-regulator and its ability to promote PTBP1-dependent exon repression (Hüttelmaier et al., 2001; Gromak et al., 2003; Rideau et al., 2006). More recent analyses have further shown that RAVER1 controls alternative-splicing programs in cancer cells through interactions with PTBPs and additional splicing regulators (Wedler et al., 2024). The recovery of a coherent spliceosomal component therefore links changes in local fractal geometry to a central molecular function of RAVER1.

C1 contained robust themes associated with miRNA-mediated gene silencing, RISC assembly and pre-miRNA processing. These annotations arose from coordinated changes involving AGO2, TNRC6B, PRKRA, XPO5, FXR1 and related proteins. RAVER1 depletion has previously been shown to alter the splicing of TNRC6-family transcripts, impair miR/RISC activity and deregulate post-transcriptional gene expression in A549 cells (Wedler et al., 2024). Notably, TNRC6B was identified here without reference to splicing: it emerged from altered network geometry in quantitative proteomes of RAVER1-KO versus parental A549 cells. The identification of a distinct miR/RISC-associated component thus independently recovered this experimentally established consequence of RAVER1 loss at the level of local network geometry.

Further components intersected additional reported functions of RAVER1. The trafficking, adhesion and cytoskeletal signals represented by C7 and C9 were consistent with the interaction of RAVER1 with vinculin, metavinculin and α-actinin and its localization at microfilament attachment sites (Hüttelmaier et al., 2001). Cell-death programmes in C6 and C9 converged with studies showing that RAVER1 regulates RIPK1 alternative splicing and thereby controls RIPK1-dependent inflammasome activation, apoptosis and pyroptotic cell death (Malireddi et al., 2023; Zhang et al., 2025). RAVER1 depletion also induces apoptosis in A549 cells (Wedler et al., 2024), and RAVER1 has been implicated in the response of AML cells to BCL2 inhibition in a CRISPR screen (Wang et al., 2023). Mitochondrial and innate immune annotations associated with C8 were also compatible with the reported interaction of RAVER1 with MDA5 and its capacity to enhance MDA5-dependent antiviral signalling (Chen et al., 2013). These correspondences do not establish that the respective component proteins are direct RAVER1 targets but demonstrate that the fractal analysis recovers multiple biological processes previously connected to RAVER1 in independent experimental settings.

In addition to these literature-supported functions, the component analysis identified several candidate roles of RAVER1 (Fig. 5a). Three gene-disjoint components converged on successive stages of tRNA biology: RNase P/MRP-associated 5′-end processing in C1, threonylcarbamoyladenosine modification in C5 and tRNA aminoacylation in C10. The recurrence of related annotations across independently connected components suggests coordinated reorganization of tRNA maturation and translational homeostasis. This association has not, to our knowledge, been linked directly to RAVER1 and therefore represents a hypothesis generated by the fractal geometry analysis.

Additional candidate functions included MICOS-associated mitochondrial membrane and cristae organization in C8, chromatin and transcriptional regulation in C4, membrane trafficking in C7 and Hedgehog signalling in C3. C6 further connected methionine salvage with apoptosome-related processes, whereas C9 contained adhesion- and cell-death-associated annotations. These modules extend the response beyond the established RNA-regulatory functions of RAVER1 and implicate broader changes in mitochondrial organization, metabolism, cellular architecture and stress adaptation. Because these signals were derived using a physical-plus-functional STRING scaffold, they were initially considered functionally informed predictions rather than annotation-independent evidence of new RAVER1 mechanisms.

The prediction of altered Hedgehog signalling was of particular interest because vertebrate Hedgehog signal transduction is closely coupled to the primary cilium (Goetz and Anderson, 2010), whereas RAVER1 has not previously been implicated in ciliogenesis. We therefore tested whether the network-derived prediction was accompanied by a ciliary phenotype using siRNA-mediated RAVER1 depletion in A549 cells and quantified ciliation (Fig. 5b–d and Fig. S2). C3 (KIF7, SUFU, GAN) and C7 (RAB10, RILPL1, RILPL2) contain components of ciliary and Hedgehog signalling. RAVER1 depletion increased the fraction cells with ARL13B- and acetylated tubulin-positive primary cilia by 2.57-fold compared to controls (control normalized to 100%; p < 0.01) and lengthened cilia (4.719 ± 1.406 µm versus 3.542 ± 0.853 µm in controls; p < 0.01). These findings support a role for RAVER1 in limiting ciliation and cilium length under the tested A549 culture conditions, a function not described before.

Preranked GSEA (Subramanian et al., 2005) using limma moderated t statistics (Smyth, 2004) without a fold-change cutoff detected no significant enrichment among the tested cilia, Hedgehog, mRNA-processing, splicing, microRNA or gene-silencing focus sets (adj. p > 0.05; Supplementary Table S1). Their identification within DIFFRACT components therefore supports the ability of network geometry to reveal functional reorganization beyond the coordinated abundance shifts detected by conventional gene-set enrichment.

To assess the recurrence of functional findings in a second cellular model, we applied DIFFRACT to RAVER1-KO and WT HEK293T proteomes. Across the evaluated scaffolds and weighting methods, 52 of the 338 distinct terms enriched in the A549 C1–C10 components were also significantly enriched in HEK293T, including cilium assembly and organization, tRNA metabolism and mitochondrial cristae formation (Supplementary Table S2). In contrast, mRNA processing and splicing were significantly detected by abundance-ranked GSEA in HEK293T but not by DIFFRACT component enrichment, indicating cell-context-dependent representation of these processes in abundance changes and network reorganization.

**Figure 5.**
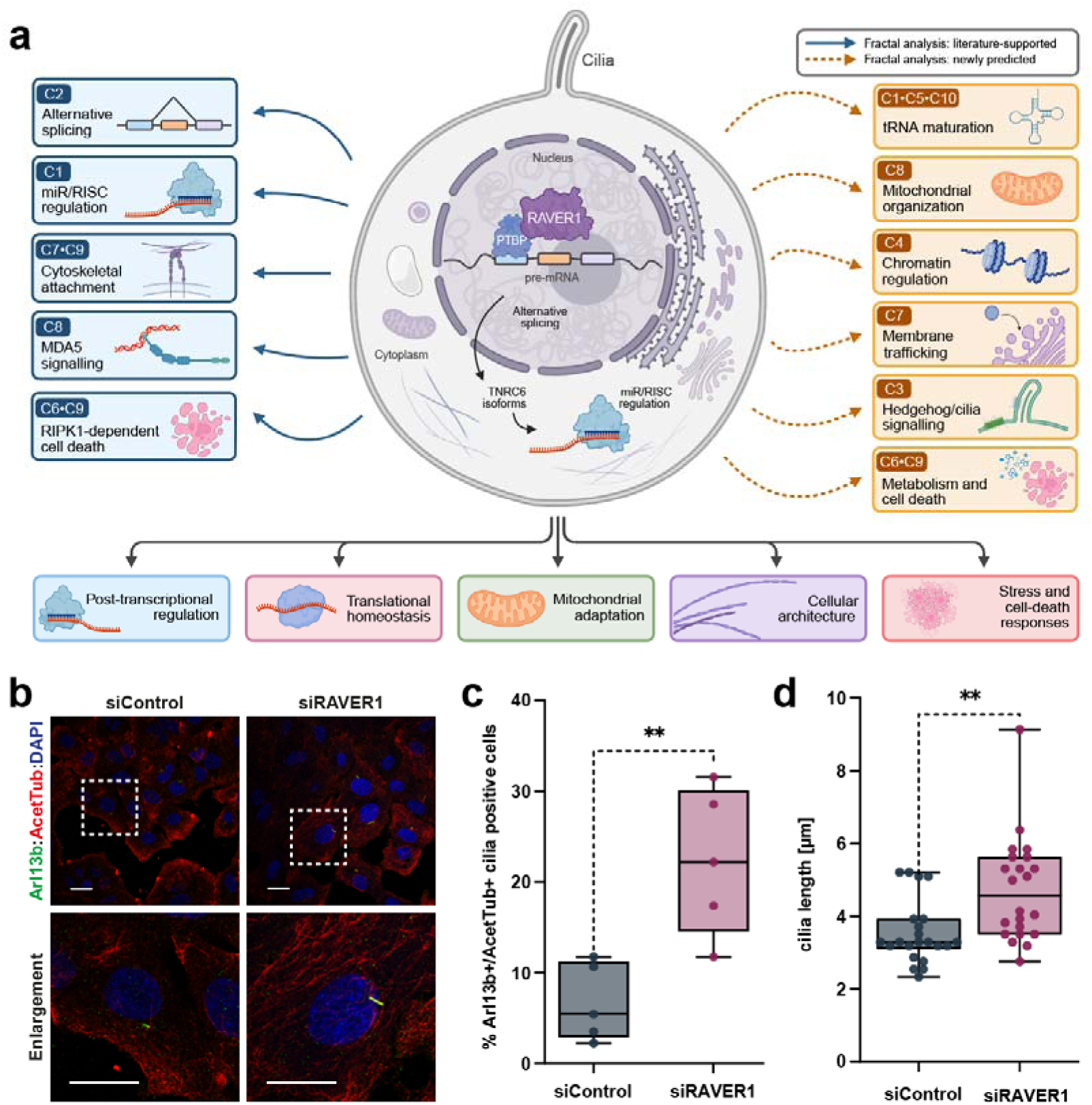
Fractal network geometry recovers established RAVER1 functions and predicts a role in ciliogenesis. **a)** Integrated model of the functional programs associated with RAVER1 deletion. Changes in local fractal geometry within MI-weighted components of the STRING physical-plus-functional network were consolidated into literature-supported and newly predicted functions. The central cell depicts the established RAVER1–PTBP axis connecting alternative splicing with the regulation of TNRC6 isoforms and miR/RISC activity. Solid blue arrows indicate functions recovered by the fractal analysis that are supported by previous experimental studies of RAVER1: alternative splicing (C2), miR/RISC regulation (C1), cytoskeletal attachment and cellular adhesion (C7 and C9), MDA5-mediated signalling (C8), and RIPK1-dependent cell-death regulation (C6 and C9). Dashed ochre arrows indicate functions newly proposed by the fractal-network analysis: tRNA maturation (C1, C5 and C10), mitochondrial organization (C8), chromatin regulation (C4), membrane trafficking (C7), Hedgehog signaling (C3), and metabolic and cell-death programmed (C6 and C9). The lower panel summarizes the broader cellular processes in which these modules converge, including post-transcriptional regulation, translational homeostasis, mitochondrial adaptation, cellular architecture, and stress and cell-death responses. Component labels refer to the MI-derived connected components identified after mapping selected ΔLFD outliers onto the STRING interaction scaffold. **b)** Representative immunofluorescence images of A549 cells transfected with 15 nM control siRNA (siControl) or RAVER1-targeting siRNA (siRAVER1) and analysed 72 h after transfection. Primary cilia were detected by co-staining for ARL13B (green) and acetylated α-tubulin (AcetTub; red); nuclei were stained with DAPI (blue). Dashed boxes indicate the regions shown at higher magnification below, and white arrows indicate cilia positive for both ARL13B and acetylated α-tubulin. Scale bars, 20 µm. **c)** Percentage of ciliated cells following control or RAVER1-targeting siRNA treatment. Cilia were defined as structures positive for both ARL13B and acetylated α-tubulin, and the percentage of ciliated cells was calculated relative to the number of DAPI-stained nuclei. Points represent independent biological replicates (n = 5). **d)** Length of individual cilia positive for both ARL13B and acetylated α-tubulin in control and RAVER1-depleted cells. Box plots show the median and interquartile range. Statistical significance was assessed using a two-sided Student’s t-test; **p < 0.01. RAVER1 depletion increased both the proportion of ciliated cells and cilium length, experimentally supporting the network-derived prediction that RAVER1 contributes to the regulation of ciliogenesis.

## Discussion

DIFFRACT provides a framework for identifying perturbation-associated changes in the network context of proteins. By integrating quantitative proteomics with interaction scaffolds and measuring changes in LFD (Pu et al., 2014), it complements analyses centered on differential protein abundance. The approach builds on earlier strategies for identifying molecular subnetworks from expression data and on the development of node-specific descriptions of network geometry. It combines condition-dependent edge weighting with a local geometric readout that prioritizes proteins whose surrounding network organization changes following perturbation (Ideker et al., 2002; Xiao et al., 2021).

DIFFRACT addresses a question shared with co-expression and differential network analysis: how molecular relationships change beyond differences in individual abundances. Weighted gene co-expression network analysis (WGCNA) (Langfelder & Horvath, 2008) transforms correlations between molecular profiles into weighted adjacencies and commonly uses topological overlap to identify modules, which can subsequently be related to experimental traits. DiffCoEx (Tesson et al., 2010) extends this framework by grouping genes according to changes in their correlation patterns between conditions, including changes within and between modules. Thus, detecting functional responses that are not apparent from differential abundance is an established objective of network analysis. DIFFRACT, however, contributes a geometric description of these responses: molecular associations are translated into effective distances on a supplied interaction scaffold, and changes are summarized by the growth of neighborhoods around individual proteins. Connected components are then extracted from selected ΔLFD outliers, rather than defined by clustering co-expression similarity.

Mutual information provides a flexible basis for quantifying molecular associations. In developing ARACNE, Margolin et al. (2006) showed that inference based on mutual-information rankings can be relatively resilient to estimation errors, supporting its use in network analysis without establishing the reliability of individual estimates in the present sampling regime. Approaches such as ARACNE and the more recent integrative differential dependency networks framework, iDDN (Wang et al., 2025), infer network relationships from molecular profiles, with iDDN extending this task across multiple omics layers. A central challenge is establishing whether inferred edges represent biological relationships, particularly in systems where an experimentally established reference network is unavailable. DIFFRACT avoids reconstructing connectivity *de novo* by assigning condition-specific distances to edges supported by an existing interaction scaffold. Its geometric readout then summarizes how these distances collectively alter neighborhood growth around each protein, capturing responses that extend beyond individual edges and immediate neighbors. This focuses inference on changes in network organization while retaining dependence on the coverage and quality of the chosen scaffold.

Integrating molecular measurements with prior interaction knowledge is also central to approaches such as PANDA (Glass et al., 2013) and COSMOS (Dugourd et al., 2021). PANDA combines gene expression, transcription-factor binding motifs and protein interactions through message passing to infer regulatory networks (Glass et al., 2013). COSMOS integrates transcriptomic, phosphoproteomic and metabolomic measurements with prior signaling, regulatory and metabolic relationships to generate causally coherent mechanistic hypotheses (Dugourd et al., 2021). DIFFRACT operates at a different interpretative level: its undirected weighted geometry identifies proteins whose network neighborhoods change, without assigning regulatory direction or requiring measurements of transcription-factor or kinase activity. These geometric candidates could therefore guide subsequent investigation using more mechanistically constrained network models. The controlled edge-rescaling experiments support the sensitivity of this readout to regional changes, while its invariance to uniform rescaling establishes that it measures relative network organization.

The biological application illustrates the potential value of this geometric perspective. In A549 cells, DIFFRACT highlighted RNA-regulatory and ciliary processes that were not significantly enriched by abundance-ranked GSEA, and experimental follow-up identified increased ciliation and cilium length after RAVER1 depletion. Recurrence of selected functional annotations in HEK293T further supports investigating these processes across cellular contexts. These findings support DIFFRACT as a complementary strategy for generating experimentally testable hypotheses from proteomic data. Future comparisons on well-characterized perturbation datasets could establish when changes in LFD provide additional biological insight beyond existing network-analysis methods.

The recovery of RNA processing and post-transcriptional regulatory functions by DIFFRACT is consistent with established RAVER1 biology. RAVER1 interacts with PTBP1 and contributes to the regulation of alternative splicing, as demonstrated for α-tropomyosin transcripts (Gromak et al., 2003). More recently, RAVER1 was shown to regulate alternative splicing of TNRC6-family transcripts, including TNRC6B exon 6, whose inclusion promotes AGO2 association in protein-fragment assays (Wedler et al., 2024). These findings connect RAVER1-dependent splicing to the molecular interactions involved in miRNA-mediated regulation.

The increase in ciliation and cilia length following RAVER1 depletion in A549 cells identifies ciliogenesis as an additional phenotype associated with RAVER1 loss (Fig. 5b–d). This observation is consistent with the functional annotations of C3, containing KIF7, SUFU and GAN, and C7, containing RAB10, RILPL1 and RILPL2. KIF7 organizes the ciliary tip compartment and the distribution of Hedgehog signalling components, whereas LRRK2-dependent phosphorylation of RAB10 enhances its association with RILPL1 and inhibits ciliogenesis (He et al., 2014; Dhekne et al., 2018). The phenotype therefore supports the functional relevance of the processes highlighted by these components, without establishing which constituent proteins mediate the response. Whether RAVER1 acts through Hedgehog signaling, the LRRK2–RAB10–RILPL1 axis, or altered splicing of ciliary transcripts remains unresolved.

A potential connection to epithelial responses to infection merits investigation. CRISPRi analysis of COVID-19-associated genomic regions implicated LZTFL1 and RAVER1 as regulated genes, with the authors relating LZTFL1 to airway ciliogenesis and RAVER1 to MDA5-dependent antiviral signaling (Fink-Baldauf et al., 2022). Our findings raise the possibility that RAVER1 also influences an epithelial ciliary phenotype. However, primary cilium formation in A549 cells is distinct from the motile multiciliation responsible for airway mucociliary clearance, and the present experiments do not establish an effect on viral entry, clearance or disease severity. Testing RAVER1 in differentiated airway epithelial models would be required to evaluate such a connection.

The convergence of three components, C1, C5 and C10, on tRNA processing, modification and aminoacylation suggests that the consequences of RAVER1 loss extend beyond its established roles in pre-mRNA splicing and miRNA-mediated regulation to the machinery supporting translation. The KEOPS-associated signal in C5 connects this response to t^6^A modification, which contributes to accurate translational decoding (Daugeron et al., 2011). C1 further contains the RNase P/MRP-associated proteins POP1, POP4 and RPP25L, providing a potential link between tRNA maturation and ribosome biogenesis: human RNase MRP cleaves pre-rRNA, and RPP25L depletion impairs processing of its ITS1 region (Goldfarb & Cech, 2017; Dodel et al., 2024). Together, these findings raise the hypothesis that RAVER1 contributes, directly or through downstream RNA-regulatory effects, to coordinating the maturation of RNAs required for protein synthesis.

The RNA-processing findings also suggest a possible connection between nuclear organization and cytoplasmic RNA regulation. RAVER1 localizes to perinucleolar structures (Hüttelmaier et al., 2001), while the long noncoding RNA PNCTR, transcribed predominantly by RNA polymerase I from an rDNA intergenic spacer, assembles a perinucleolar compartment that sequesters PTBP1 and promotes cell survival (Yap et al., 2018). TNRC6 proteins, in turn, localize to P-bodies and participate in miRNA-mediated repression and mRNA destabilization (Lazzaretti et al., 2009). We therefore speculate that RAVER1 could connect perinucleolar responses to perturbed ribosome biogenesis with cytoplasmic RNA regulation involving P-body-associated factors.

Mitochondrial cristae organization provides another example of functional convergence without identical protein involvement. The A549 signal involved MICOS-associated proteins, whereas the corresponding HEK293T enrichment involved ATP synthase subunits. This distinction is biologically plausible because mitochondrial contact-site machinery and ATP synthase organization contribute through different mechanisms to inner-membrane architecture (Harner et al., 2011; Rabl et al., 2009). The shared annotation therefore suggests a process worth examining across models, rather than replication of a specific molecular mechanism. Ultrastructural imaging and measurements of respiratory function would determine whether the inferred changes correspond to altered cristae morphology or mitochondrial physiology.

The present implementation has several limitations. Mutual information estimated from three biological replicates using two bins provides limited resolution and is susceptible to finite-sample bias (Paninski, 2003). Larger replicate series are therefore needed to assess the stability of the resulting network patterns, while direct benchmarking should establish when the geometric readout adds information beyond existing network-analysis methods. ΔLFD outliers prioritize unusual responses relative to the network-wide distribution without controlling a protein-level false discovery rate; the resulting components should consequently be regarded as candidates for further investigation. LFD is likewise interpreted as a comparative descriptor over the evaluated neighborhood ranges, rather than evidence of universal fractal scaling.

The choice of interaction scaffold also shapes biological interpretation. Incorporating functional associations from STRING resolved components that were more readily interpretable than those obtained from physical interactions alone. However, these associations partly draw on knowledge related to the annotations used for enrichment, precluding enrichment as independent validation of the components (Szklarczyk et al., 2025). Coverage of approximately 9,000 proteins further limits access to unmeasured proteins, isoforms, post-translational modifications and compartment-specific regulation. Despite these constraints, DIFFRACT identified diverse functional associations, some recurring in HEK293T, and guided experimental identification of a ciliary phenotype in A549 cells. These findings support its utility for biological hypothesis generation, while broader generalization and the distinction between direct RAVER1 effects and adaptation to chronic knockout require further investigation.

Future studies should establish how the direction of ΔLFD relates to functional outcomes, including whether positive and negative responses distinguish enhancement from impairment in defined biological contexts. Application to RNA-seq and other molecular profiles would extend this analysis to additional regulatory layers. Targeted cellular perturbations could test the predicted relationships between distant network regions suggested by the sensitivity of LFD beyond immediate neighbors. An R package is under development to facilitate reproducible application and wider adoption. Together, these developments extend DIFFRACT as a framework for connecting molecular perturbations to changes in network organization and experimentally testable cellular functions.

## Materials and Methods

### Molecular interaction networks

Physical molecular interaction networks were obtained for *Escherichia coli*, *Arabidopsis thaliana*, *Caenorhabditis elegans* and *Homo sapiens* from BioGRID (v4.4.240) (Stark et al., 2006), IntAct (release 252, January 2026) (del Toro et al., 2022) and STRING (v12.0) (von Mering et al., 2003; Szklarczyk et al., 2025). Protein identifiers were standardized to UniProt accessions (The UniProt Consortium, 2025), self-interactions and duplicate edges were removed, and all networks were represented as undirected graphs. Analyses were restricted to the largest connected component (LCC) of each network.

The resulting BioGRID (Stark et al., 2006), IntAct (del Toro et al., 2022) and STRING (von Mering et al., 2003; Szklarczyk et al., 2025) LCCs contained, respectively, 3,429 nodes and 23,903 edges, 1,671 nodes and 2,889 edges, and 350 nodes and 3,134 edges for *E. coli*; 10,866 nodes and 51,702 edges, 7,108 nodes and 36,345 edges, and 668 nodes and 4,242 edges for *A. thaliana*; 7,650 nodes and 26,167 edges, 4,599 nodes and 11,338 edges, and 428 nodes and 1,367 edges for *C. elegans*; and 20,154 nodes and 850,733 edges, 5,611 nodes and 14,637 edges, and 7,167 nodes and 43,180 edges for *H. sapiens*.

An additional human STRING (von Mering et al., 2003; Szklarczyk et al., 2025) network containing high-confidence physical and functional associations (combined score > 0.9) was examined to increase functional coverage and permit the resolution of connected proteins into biologically interpretable modules. After mapping the quantified proteins and restricting the network to its largest connected component, the resulting WT and RAVER1-KO (see below) networks contained 6,056 proteins connected by 56,343 edges and 6,068 proteins connected by 56,694 edges, respectively. Because functional STRING associations incorporate annotation-dependent evidence, this expanded network was used for biological interpretation and module resolution rather than as an independent validation of functional enrichment.

### Global box-covering analysis

Global fractal scaling was evaluated using the network box-covering framework introduced by Song, Havlin and Makse (2005) and its randomized covering formulation (Song et al., 2007). For each graph, the number of radius-defined boxes required to cover the network was approximated using a randomized sequential ball-covering algorithm. At each iteration, an uncovered node was selected randomly as the center of a box, and all nodes within a shortest-path radius l_B_ of that center were assigned to the box. This procedure was repeated until every node was covered.

Because the resulting cover depends on the order in which box centers are selected, 100 independent covering trials were performed for each value of l_B_. The mean number of boxes across trials was denoted by N_B_(l_B_). This represents a heuristic approximation of the minimum network cover, as exact minimum box covering is computationally intractable for large graphs.

For a fractal network, the number of boxes is expected to follow:

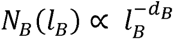

where d_B_ is the box-counting fractal dimension.

After normalization by the total number of nodes N, the relationship becomes:

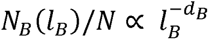

The box-counting fractal dimension d_B_ was estimated as the negative slope of the approximately linear relationship between log_10_[N_B_(l_B_)/N] and log_10_(l_B_). Candidate scaling intervals were required to contain at least five consecutive radii, to exclude non-finite and saturated values with N_B_(l_B_) ≤ 1, and to exhibit a negative fitted slope. The interval with the highest adjusted R^2^ was identified first. Among intervals within 0.01 of this maximum, the interval containing the greatest number of radii was selected, with the lowest root-mean-square error (RMSE) used to resolve remaining ties. This procedure restricted dimension estimation to the scaling region preceding the finite-size plateau.

### Multifractal network analysis

Heterogeneity in network scaling was evaluated using a node-centred box-growing procedure based on the node-based multifractal analysis framework described by Xiao, Chen and Bogdan (2021).

Broad, non-degenerate f(α) spectra were interpreted as evidence that different regions of a network exhibit distinct local scaling behaviour. Spectrum position and width were treated as descriptive properties and compared across organisms and interaction resources.

### Degree-preserving network controls

To determine whether the observed scaling properties could be explained by network size and degree distribution alone, degree-preserving null networks were generated by repeated edge switching. In each switch, two edges were exchanged while preserving the degree of every node and avoiding self-loops and duplicate edges.

Null networks were generated independently for each empirical interaction network and subjected to the same global box covering, multifractal and local fractal dimension analyses. Empirical results were compared with the corresponding null distributions using the same fitting and quality control criteria.

### Local fractal dimension

Node-wise local fractal dimension (LFD) was calculated using a box-growing procedure adapted from the node-based fractal dimension introduced by Xiao, Chen and Bogdan (2021). Whereas global box covering assigns a single scaling exponent to the complete graph, LFD quantifies the rate at which the neighbourhood surrounding an individual node expands over increasing shortest-path radii.

For node i, M_i_(r) denotes the cumulative number of other nodes reachable within shortest-path radius r. The focal node itself was excluded from this count. Local neighbourhood growth was represented as:

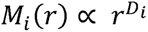

where D_i_ is the local fractal dimension of node i. Equivalently:

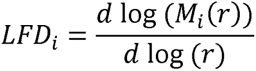

In practice, LFD_i_ was estimated as the ordinary least-squares slope of log[M_i_(r)] against log(r) across the complete set of unique, finite and positive shortest-path radii at which neighborhood growth occurred. Nodes for which an adequate growth curve could not be fitted were excluded from the corresponding comparison.

For an unweighted network, each edge was assigned unit distance. The resulting value represented the structural LFD inherent to the architecture and topology of the network, LFD_structural_. For a proteome-weighted network, shortest paths were calculated using proteome-derived edge distances. The resulting value represented total weighted LFD, LFD_total_(i, c), where c denotes the biological condition.

Total LFD was decomposed into structural and dynamic, proteome-dependent components:

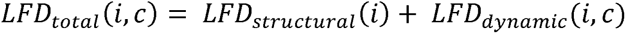

This decomposition expresses weighted LFD relative to the unweighted structural baseline. However, the resulting dynamic component remains conditional on the interaction scaffold.

### Controlled edge-rescaling analysis

The response of LFD to spatially localized changes in edge distance was evaluated by systematically rescaling selected edges in the empirical interaction networks. Edges located inside, adjacent to or outside a predefined perturbation region were multiplied by a rescaling factor s:

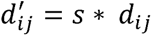

where d_ij_ is the original edge distance and d′_ij_ is the rescaled distance. Values of s < 1 represented local contraction, s = 1 represented the unperturbed network and s > 1 represented local relaxation.

Following each perturbation, LFD was recalculated for every evaluable node. The relationship between the imposed edge-rescaling factor and the resulting LFD response was evaluated using the adjusted coefficient of determination and the root-mean-square error of the node-wise scaling fits.

Spatial localization of the response was evaluated from the correlation between dynamic LFD values of adjacent nodes, the mean absolute dynamic-LFD difference between adjacent nodes and the shortest-path distance over which the response propagated from the perturbed region.

### siRNA knockdown and immunofluorescence microscopy

A549 cells were cultured in DMEM supplemented with 10% fetal bovine serum (FBS) at 37 °C and 5% CO_2_. For siRNA-mediated knockdown, 7 × 10^5^ cells were transfected with 15 nM non-targeting control siRNA (siControl: UUUGUACUACACAAAAGUACUG) or RAVER1-targeting siRNA pool (siRAVER1: CUCGCUUGUUGAAGAUGAA; CCAGCACUCCAUGAAGAUA; AUGUCCGACUCCCACAGAC; AUAGGCCAGAUGGGCAGGC; UAAGGCUGCAGGGUUUCAG) using Lipofectamine RNAiMAX (Thermo Fisher Scientific) according to the manufacturer’s instructions (Supplementary Figure S2). Cells were seeded onto coverslips 24 h after transfection. At 72 h after transfection, cells were fixed with paraformaldehyde and processed for immunofluorescence staining as described in Wedler et al. (2024). Primary antibodies against ARL13B (Proteintech, 17711-1-AP) and acetylated α-tubulin (Sigma, T7451) were used to label primary cilia, and nuclei were counterstained with DAPI. As secondary antibodies Cy™3 AffiniPure® F(ab’)_2_ Fragment Donkey Anti-Mouse IgG (H+L) (Dianova, 715-166-150) and Alexa Fluor® 488 AffiniPure® F(ab’)_2_ Fragment Donkey Anti-Rabbit IgG (H+L) (Dianova, 711-546-152) were used. Images were acquired on a Leica SP8X confocal microscope equipped with a white-light laser, HyD detectors and a 63× oil-immersion objective using sequential acquisition.

Images were quantified using Fiji (version 1.54). The total number of cells was determined by counting DAPI-stained nuclei. Cilia were identified as structures positive for both ARL13B and acetylated α-tubulin. The number of double-positive cilia was divided by the number of nuclei to calculate the percentage of cilia positive cell fraction for each of five biological replicates. Cilium length was measured along individual structures positive for both markers.

### Cell culture and generation of RAVER1-KO cells

WT and RAVER1-KO A549 human lung adenocarcinoma and HEK293T human embryonal kidney cells were analyzed using three biological replicates per condition. Cells were grown to confluence in 15-cm culture dishes, washed with phosphate-buffered saline (PBS), detached by scraping and collected by centrifugation at 500 × g for 5 min. The resulting cell pellets were processed for proteomic analysis. Generation and validation of the RAVER1-KO cells were performed as described by Wedler et al. (2024). WT and KO samples were processed and measured using the same proteomic workflow.

### Proteomic sample preparation and LC-MS analysis

Samples were prepared using a modified filter-aided sample preparation (FASP) protocol based on Wiśniewski et al. (2009). After tryptic digestion, peptides were analyzed on an Ultimate 3000 RSLC nano-HPLC system coupled to a timsTOF Ultra-2 mass spectrometer with a CaptiveSpray ion source (Bruker Daltonik). Peptides were loaded onto a C18 trap column (Acclaim PepMap, 300 µm × 5 mm, 5 µm, 100 Å; Thermo Fisher Scientific), washed with 0.1% trifluoroacetic acid, and separated on a self-packed Picofrit C18 nanospray column (75 µm × 250 mm, 10 µm tip ID; New Objective) packed with ReproSil-Pur C18 (1.9 µm, 120 Å; Dr. Maisch). Elution used a 3–30% acetonitrile gradient in 0.1% formic acid over 60 min at 300 nL/min.

Data were acquired using DIA-PASEF (data independent acquisition - parallel acquisition serial fragmentation) with an MS *m/z* range of 100–1700, an ion mobility range of 0.64–1.45 V s/cm^2^, and a 100-ms ramp time. Each cycle comprised eight MS/MS ramps with 24 windows spanning m/z 400– 1000 and 0.64–1.37 V s/cm^2^. Collision energy increased linearly from 20 eV at 0.6 V s/cm² to 59 eV at 1.6 V s/cm^2^.

DIA data were searched in real time using directDIA in Spectronaut (version 19; Biognosys) with Proteoscape (Bruker Daltonik) against the UniProt (The UniProt Consortium, 2025) human database (February 2025; 20,427 entries). Default identification settings were used: precursor q-value ≤0.01 and posterior error probability (PEP) ≤0.2; protein q-value ≤0.01 and PEP ≤0.75; and at least two peptides per protein. Searches allowed two missed cleavages and peptides of at least seven amino acids. Methionine oxidation and protein N-terminal acetylation were variable modifications; cysteine carbamidomethylation was fixed. Only proteotypic peptides were used for protein quantification.

### Proteomic data processing

Protein abundances were assembled into a protein-by-sample matrix. Protein identifiers were standardized before mapping to the interaction networks. Zero and missing observations were retained as explicit measurement masks during the calculation of expression-derived edge metrics.

Principal component analysis was performed on the processed protein-abundance matrix to assess the overall similarity of biological replicates and separation between WT and RAVER1-KO samples.

### Differential protein-abundance analysis

Differential protein abundance between RAVER1-KO and WT cells was evaluated for 8,966 quantified proteins. Resulting p-values were corrected for multiple testing using the Benjamini-Hochberg (1995) procedure. Proteins were classified as significantly differentially abundant at adj. p-value < 0.05 and |log_2_(FC)| > 1.

### Construction of proteome-weighted interaction networks

Because the proteomic measurements were obtained from human A549 cells, expression-weighted analyses were restricted to the human interaction networks. Quantified proteins were mapped to the BioGRID (Stark et al., 2006), IntAct (del Toro et al., 2022) and STRING (von Mering et al., 2003; Szklarczyk et al., 2025) scaffolds. Network topology was held fixed within each comparison, and proteomic measurements were used only to assign edge distances.

After restriction to proteins that could be evaluated from the WT A549 proteomes, the BioGRID LCC contained 7,809 nodes and 422,036 edges, the IntAct LCC contained 3,338 nodes and 8,282 edges, and the physical STRING LCC contained 1,381 nodes and 6,605 edges. For direct comparison of WT and RAVER1-KO conditions, restriction to proteins evaluable in both conditions resulted in an LCC of 7,571 nodes and 407,398 edges for BioGRID, 3,223 nodes and 7,918 edges for IntAct, and 1,363 nodes and 6,437 edges for physical STRING. In the extended STRING physical-plus-functional analysis, the condition-specific LCCs contained 6,056 nodes and 56,343 edges in the WT network and 6,068 nodes and 56,694 edges in the RAVER1-KO network.

### Proteome-derived edge associations and distances

Associations between the abundance profiles of interacting proteins were quantified using mutual information (MI), absolute Pearson correlation, distance correlation (dCor), and a variance/missingness-based metric, termed VarMiss, deriving edge distances from marginal protein variance and pairwise data completeness without using edge-specific dependence information.

### Expression-permutation controls

Expression-permuted networks were generated to determine whether proteome-weighted LFD retained information beyond the marginal distributions and observation patterns of individual proteins. For each protein, abundance values were independently permuted across samples while preserving its zero and missing-value masks. For each permutation, edge associations, effective distances, weighted shortest paths and node-wise LFD values were recalculated using the same workflow applied to the physiological proteomes.

### Evaluation of edge-weighting strategies

The four weighting strategies were evaluated according to their ability to translate proteomic coordination into reproducible changes in local network geometry. Evaluation criteria included the separation of empirical and expression-permuted edge-weight distributions, the shift in total LFD relative to structural and permutation controls, the quality of node-wise scaling fits, spatial coherence between adjacent nodes and discrimination between empirical and permuted proteomic profiles.

For every interaction resource and weighting method, scaling quality was summarized using the median adjusted R^2^ and median LFD-fit RMSE across evaluable nodes. Spatial coherence was assessed by the correlation of dynamic LFD between adjacent nodes:

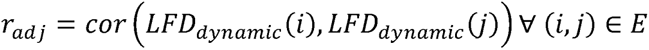

Null discrimination was calculated as an oriented summary of the separation between empirical and permutation-control results, with the direction defined such that larger positive values indicated stronger discrimination in the expected direction.

MI was selected as the primary edge-weighting metric because it showed the most consistent combined performance across interaction resources. Pearson correlation, dCor and VarMiss were retained as complementary sensitivity analyses rather than treated as independent biological replicates.

### Condition-dependent changes in local fractal geometry

Proteome-derived edge distances were calculated independently for WT and RAVER1-KO A549 cells. For each condition, total LFD was decomposed into structural and dynamic components (see Materials and Methods – Local Fracal Dimension). The response to RAVER1 depletion was defined as the difference in dynamic LFD between KO and WT cells.

The decomposed formulation was nevertheless retained conceptually because it distinguishes proteome-dependent reorganization from the fixed structural contribution. Positive and negative ΔLFD values represent opposite directions of change in local neighborhood expansion.

For each interaction scaffold and weighting method, node-wise ΔLFD values were standardized relative to their distribution across evaluable nodes:

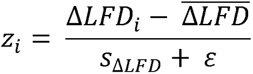

where 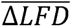 and 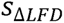 denote the mean and sample standard deviation across nodes, respectively, and ε is a numerical stabilizer. These standardized scores quantify how unusual a node’s response is relative to the network-wide distribution; they do not represent test statistics calibrated against a node-specific experimental null distribution. Nodes with 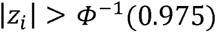, approximately 1.96, were selected as network-relative ΔLFD outliers.

### Identification of connected components among ΔLFD outliers

Nodes selected as network-relative ΔLFD outliers were mapped onto their respective interaction scaffold, and the induced subgraph was decomposed into connected components. Components containing at least five proteins were retained, using five proteins as a pragmatic heuristic threshold to exclude isolated nodes and very small subnetworks while preserving components of sufficient size for the subsequent functional overrepresentation analysis (see below). This cutoff was not statistically optimized, and alternative minimum component sizes could in principle have been selected.

### Functional overrepresentation analysis and consolidation of enriched terms

Functional overrepresentation analysis was performed separately for each retained connected component using Gene Ontology (Ashburner et al., 2000; The Gene Ontology Consortium, 2023) biological process and Reactome (Milacic et al., 2024) annotations.

Significantly enriched Gene Ontology (Ashburner et al., 2000; The Gene Ontology Consortium, 2023) biological process and Reactome (Milacic et al., 2024) terms were consolidated into non-redundant component-specific themes according to overlap among their contributing proteins.

For two enriched terms A and B, overlap was quantified using the Jaccard similarity. Terms were initially grouped by average-linkage clustering of their contributing-protein sets. Groups were subsequently merged when the Jaccard similarity between their combined protein sets was at least 0.5.

### Leave-one-protein-out sensitivity analysis

The dependence of each enriched theme on individual component proteins was evaluated by repeating the overrepresentation analysis after removing one component protein at a time. The annotation universe, term sizes and original multiple-testing family were held constant across all leave-one-protein-out iterations. Terms with no remaining component hits were retained in the testing family. All term-level p-values were then subjected to Benjamini-Hochberg (1995) correction within the original component-by-annotation-collection testing family. A theme was classified as robust when it retained at least one significantly enriched constituent term after the removal of every individual component protein.

### Statistical analysis and reproducibility

All statistical analyses were performed in R. Unless specified otherwise, tests were two-sided. Multiple-testing correction was performed using the Benjamini-Hochberg (1995) procedure. Adjusted P < 0.05 was used as the significance threshold for differential abundance and functional enrichment.

Network topology was kept fixed within comparisons intended to isolate the effect of proteome-derived edge distances. The same preprocessing, shortest-path, LFD-fitting and quality-control procedures were applied to empirical, control and perturbed networks. ΔLFD-based node selection used the network-relative outlier criterion described above and was not treated as hypothesis testing; significance thresholds for differential abundance and functional enrichment were applied separately.

## Acknowledgements

This work was supported by the Deutsche Forschungsgemeinschaft (DFG) (RTG 2467, project number 391498659 “Intrinsically Disordered Proteins-Molecular Principles, Cellular Functions, and Diseases”, CRC 1664, project number 514901783 “SNP2Prot - Plant Proteoform Diversity”, and CRC 1423, project number 421152132 “Structural Dynamics of GPCR Activation and Signaling”). AS and SH acknowledge additional financial support by the DFG (RTG 2751, project number 449501615 “InCuPanC”, RU 5433, project number 468534282 “RNA in Focus”). Additionally, AS acknowledges financial support by the DFG (INST 271/404-1 FUGG; INST 271/405-1 FUGG, INST271/528-1 FUGG), the European Regional Development Funds for Saxony-Anhalt (grant numbers EFRE ZS/2024/01/183756 and ZS/2024/01/183619), and the Martin Luther University Halle-Wittenberg (Center for Structural Mass Spectrometry).

## Author Contributions

F.H. conceived and designed the study, performed data analysis and visualization, and wrote the manuscript (first draft and revision). Cell culture experiments and immunoflourescence were performed by A.W.G., T.F., and M.L. Sample preparation, mass spectrometry measurements and MS data processing were performed by D.T and C.I. O.S., R.K., S.W, and S.H. contributed to data analysis and visualization. S. H. and A.S. critically reviewed the manuscript, supervised the study, and acquired funding.

---

**Supplementary Fig. S1.**
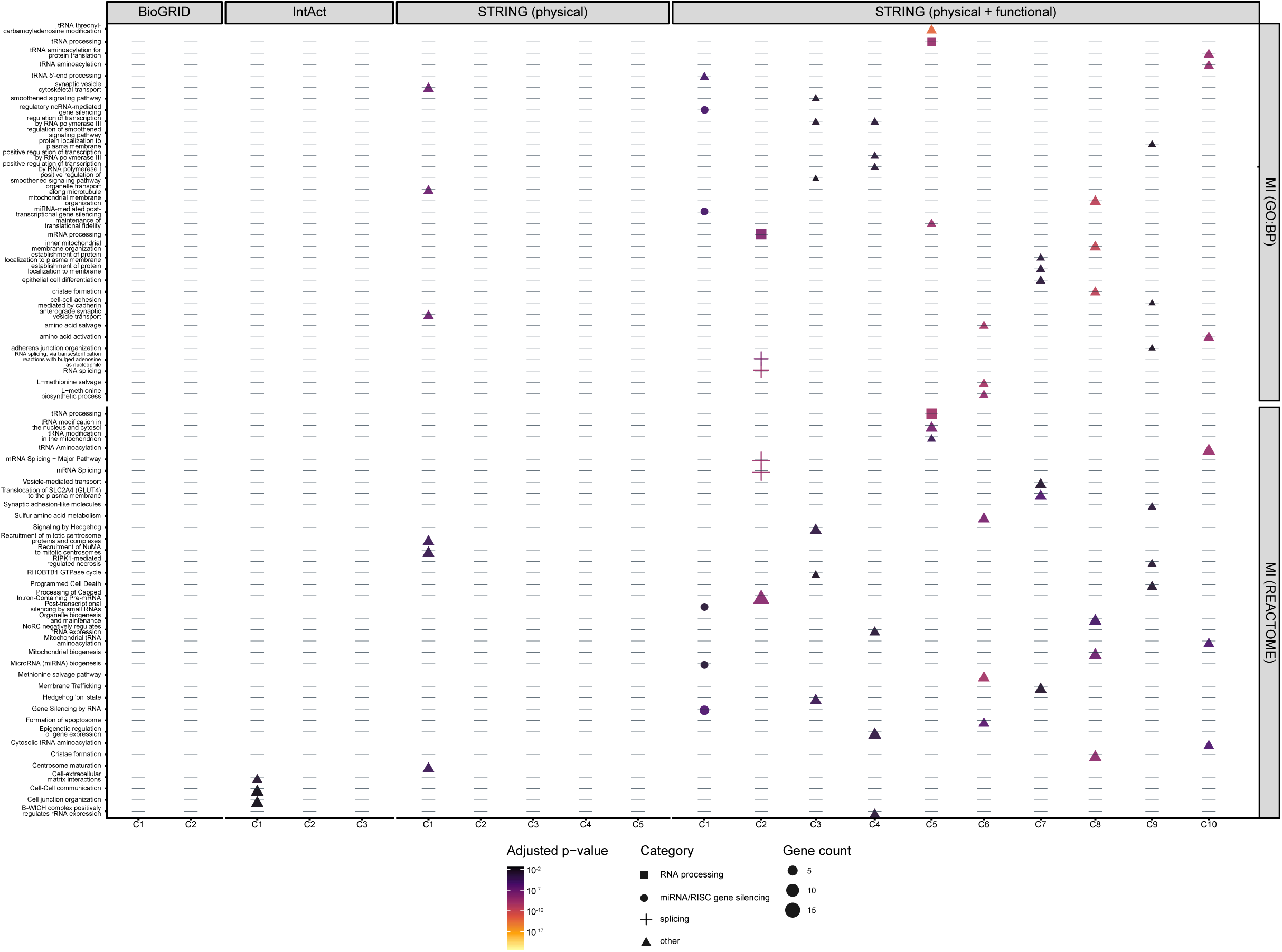
Functional enrichment of MI-derived ΔLFD components against Gene Ontology Biological Process (GO) and Reactome gene sets. The three highest-ranked significant terms are shown for each component and collection. Point colour denotes the adjusted p-value, point size the number of contributing genes and point shape prespecified functional categories related to RNA processing, miRNA/RISC-mediated gene silencing and splicing; all remaining terms are classified as other. Grey dashes denote component–term combinations without a displayed significant enrichment. C1 to C10 indicate connected components ordered by decreasing size.

**Supplementary Fig. S2.**
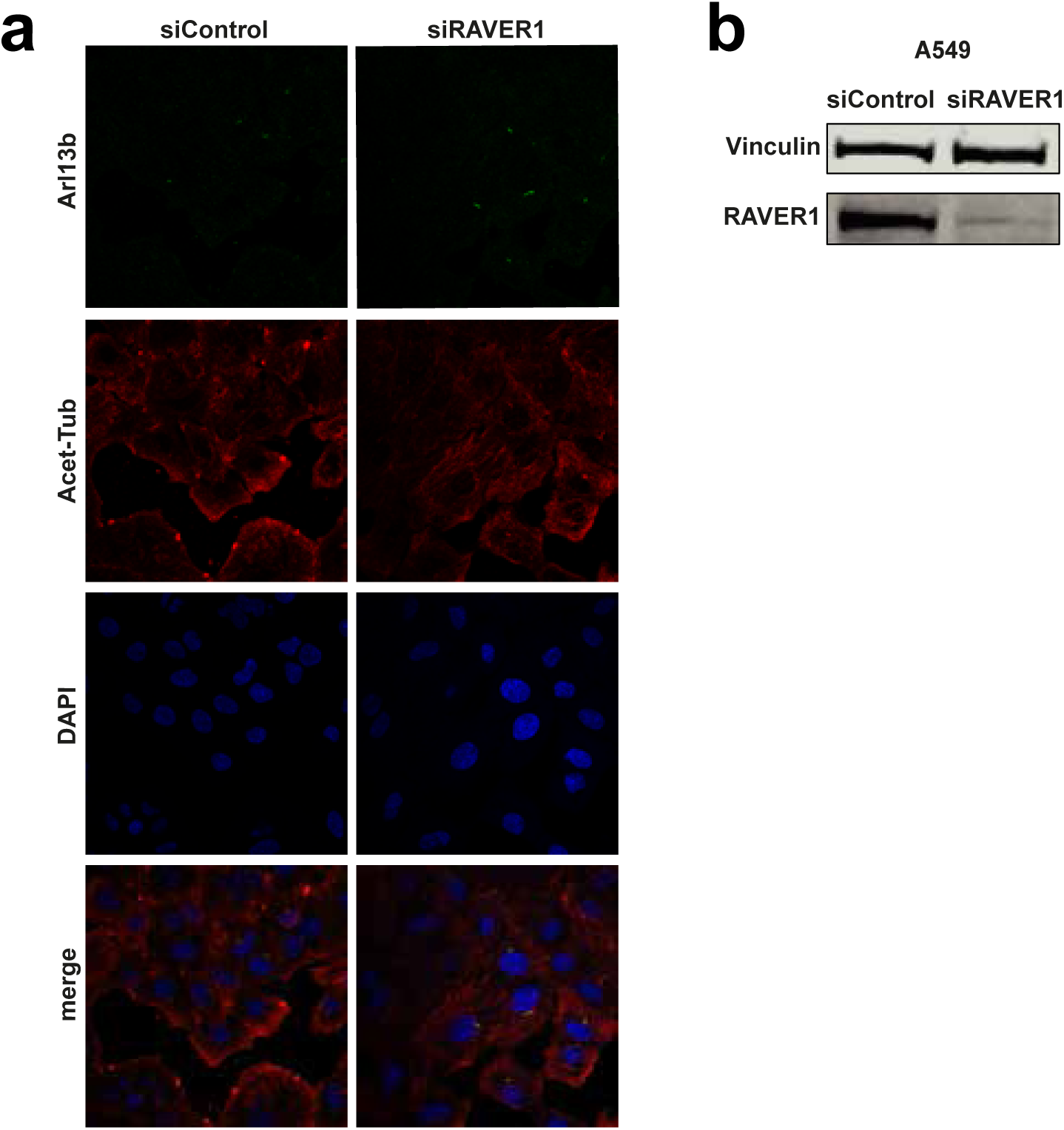
Individual fluorescence channels of control and RAVER1-depleted A549 cells. **a)** Representative confocal immunofluorescence images of A549 cells transfected with 15 nM control siRNA (siControl) or RAVER1-targeting siRNA (siRAVER1) and fixed 72 h after transfection. Individual channels show ARL13B (green), acetylated α-tubulin (red) and DAPI-stained nuclei (blue), with the corresponding merged images shown in the bottom row. Images are representative of five biological replicates. Scale bars, 20 μm. **b)** Representative immunoblot showing RAVER1 protein levels following control or RAVER1 siRNA transfection. Immunoblotting was performed as described in Wedler et al. (2024). RAVER1 was detected using an anti-RAVER1 antibody (clone 7D3, BSBS antibody facility), and vinculin was detected using an anti-vinculin antibody (Sigma, V9131) as a loading control. Immunoblot analysis was performed for each of the five biological replicates used for the immunofluorescence experiments in a).

## References

Ashburner, M. et al. Gene ontology: tool for the unification of biology. Nat. Genet. 25, 25–29 (2000).

Barabási, A.-L. & Oltvai, Z. N. Network biology: understanding the cell’s functional organization. Nat. Rev. Genet. 5, 101–113 (2004).

Benjamini, Y. & Hochberg, Y. Controlling the false discovery rate: a practical and powerful approach to multiple testing. J. R. Stat. Soc. B 57, 289–300 (1995).

Chen, H. et al. RAVER1 is a coactivator of MDA5-mediated cellular antiviral response. J. Mol. Cell Biol. 5, 111–119 (2013).

Daugeron, M.-C. et al. Gcn4 misregulation reveals a direct role for the evolutionary conserved EKC/KEOPS in the t6A modification of tRNAs. Nucleic Acids Res. 39, 6148–6160 (2011).

del Toro, N. et al. The IntAct database: efficient access to fine-grained molecular interaction data. Nucleic Acids Res. 50, D648–D653 (2022).

Dhekne, H. S. et al. A pathway for Parkinson’s disease LRRK2 kinase to block primary cilia and Sonic hedgehog signaling in the brain. eLife 7, e40202 (2018).

Dodel, M. et al. TREX reveals proteins that bind to specific RNA regions in living cells. Nat. Methods 21, 423–434 (2024).

Draghici, S., Khatri, P., Martins, R. P., Ostermeier, G. C. & Krawetz, S. A. Global functional profiling of gene expression. Genomics 81, 98–104 (2003).

Dugourd, A. et al. Causal integration of multi-omics data with prior knowledge to generate mechanistic hypotheses. Mol. Syst. Biol. 17, e9730 (2021).

Fink-Baldauf, I. M., Stuart, W. D., Brewington, J. J., Guo, M. & Maeda, Y. CRISPRi links COVID-19 GWAS loci to LZTFL1 and RAVER1. EBioMedicine 75, 103806 (2022).

Furuya, S. & Yakubo, K. Multifractality of complex networks. *Phys*. Rev. E 84, 036118 (2011).

Glass, K., Huttenhower, C., Quackenbush, J. & Yuan, G.-C. Passing messages between biological networks to refine predicted interactions. PLoS ONE 8, e64832 (2013).

Goetz, S. C. & Anderson, K. V. The primary cilium: a signalling centre during vertebrate development. Nat. Rev. Genet. 11, 331–344 (2010).

Goldfarb, K. C. & Cech, T. R. Targeted CRISPR disruption reveals a role for RNase MRP RNA in human preribosomal RNA processing. Genes Dev. 31, 59–71 (2017).

Gromak, N. et al. The PTB interacting protein raver1 regulates α-tropomyosin alternative splicing. EMBO J. 22, 6356–6364 (2003).

Harner, M. et al. The mitochondrial contact site complex, a determinant of mitochondrial architecture. EMBO J. 30, 4356–4370 (2011).

He, M. et al. The kinesin-4 protein Kif7 regulates mammalian Hedgehog signalling by organizing the cilium tip compartment. Nat. Cell Biol. 16, 663–672 (2014).

Hüttelmaier, S. et al. Raver1, a dual compartment protein, is a ligand for PTB/hnRNPI and microfilament attachment proteins. J. Cell Biol. 155, 775–786 (2001).

Ideker, T., Ozier, O., Schwikowski, B. & Siegel, A. F. Discovering regulatory and signalling circuits in molecular interaction networks. Bioinformatics 18, S233–S240 (2002).

Joyce, A. R. & Palsson, B. Ø. The model organism as a system: integrating ‘omics’ data sets. Nat. Rev. Mol. Cell Biol. 7, 198–210 (2006).

Kitano, H. Systems biology: a brief overview. Science 295, 1662–1664 (2002).

Langfelder, P. & Horvath, S. WGCNA: an R package for weighted correlation network analysis. BMC Bioinformatics 9, 559 (2008).

Lazzaretti, D., Tournier, I. & Izaurralde, E. The C-terminal domains of human TNRC6A, TNRC6B, and TNRC6C silence bound transcripts independently of Argonaute proteins. RNA 15, 1059–1066 (2009).

Malireddi, R. K. S. et al. Whole-genome CRISPR screen identifies RAVER1 as a key regulator of RIPK1-mediated inflammatory cell death, PANoptosis. iScience 26, 106938 (2023).

Margolin, A. A. et al. ARACNE: an algorithm for the reconstruction of gene regulatory networks in a mammalian cellular context. BMC Bioinformatics 7, S7 (2006).

Milacic, M. et al. The Reactome pathway knowledgebase 2024. Nucleic Acids Res. 52, D672–D678 (2024).

Mitra, K., Carvunis, A.-R., Ramesh, S. K. & Ideker, T. Integrative approaches for finding modular structure in biological networks. Nat. Rev. Genet. 14, 719–732 (2013).

Paninski, L. Estimation of entropy and mutual information. Neural Comput. 15, 1191–1253 (2003).

Pu, J., Chen, X., Wei, D., Liu, Q. & Deng, Y. Identifying influential nodes based on local dimension. EPL 107, 10010 (2014).

Rabl, R. et al. Formation of cristae and crista junctions in mitochondria depends on antagonism between Fcj1 and Su e/g. J. Cell Biol. 185, 1047–1063 (2009).

Rideau, A. P. et al. A peptide motif in Raver1 mediates splicing repression by interaction with the PTB RRM2 domain. Nat. Struct. Mol. Biol. 13, 839– 848 (2006).

Smyth, G. K. Linear models and empirical Bayes methods for assessing differential expression in microarray experiments. Stat. Appl. Genet. Mol. Biol. 3, Article 3 (2004).

Song, C., Gallos, L. K., Havlin, S. & Makse, H. A. How to calculate the fractal dimension of a complex network: the box covering algorithm. J. Stat. Mech. Theory Exp. 2007, P03006 (2007).

Song, C., Havlin, S. & Makse, H. A. Self-similarity of complex networks. Nature 433, 392–395 (2005).

Stark, C. et al. BioGRID: a general repository for interaction datasets. Nucleic Acids Res. 34, D535–D539 (2006).

Subramanian, A., et al. Gene set enrichment analysis: a knowledge-based approach for interpreting genome-wide expression profiles. Proc. Natl Acad. Sci. USA 102, 15545–15550 (2005).

Szklarczyk, D. et al. The STRING database in 2025: protein networks with directionality of regulation. Nucleic Acids Res. 53, D730–D737 (2025).

Tesson, B. M., Breitling, R. & Jansen, R. C. DiffCoEx: a simple and sensitive method to find differentially coexpressed gene modules. BMC Bioinformatics 11, 497 (2010).

The Gene Ontology Consortium. The Gene Ontology knowledgebase in 2023. Genetics 224, iyad031 (2023).

The UniProt Consortium. UniProt: the Universal Protein Knowledgebase in 2025. Nucleic Acids Res. 53, D609–D617 (2025).

von Mering, C. et al. STRING: a database of predicted functional associations between proteins. Nucleic Acids Res. 31, 258–261 (2003).

Wang, E. et al. Modulation of RNA splicing enhances response to BCL2 inhibition in leukemia. Cancer Cell 41, 164–180.e8 (2023).

Wang, Y., et al. iDDN: determining trans-omics network structure and rewiring with integrative differential dependency networks. Bioinform. Adv. 5, vbaf086 (2025).

Wedler, A. et al. RAVER1 hinders lethal EMT and modulates miR/RISC activity by the control of alternative splicing. Nucleic Acids Res. 52, 3971–3988 (2024).

Wiśniewski, J. R., Zougman, A., Nagaraj, N. & Mann, M. Universal sample preparation method for proteome analysis. Nat. Methods 6, 359–362 (2009).

Xiao, X., Chen, H. & Bogdan, P. Deciphering the generating rules and functionalities of complex networks. Sci. Rep. 11, 22964 (2021).

Yap, K. et al. A short tandem repeat-enriched RNA assembles a nuclear compartment to control alternative splicing and promote cell survival. Mol. Cell 72, 525–540.e13 (2018).

Zhang, B., et al. Raver1 links Ripk1 RNA splicing to caspase-8-mediated pyroptotic cell death, inflammation, and pathogen resistance. Proc. Natl Acad. Sci. USA 122, e2420802122 (2025).

