## Supplementary Table S2 for "Local fractal dimension as an indicator of functional reorganization in molecular interaction networks"

| collection | term_id | term | A549_compon | A549_min_adj | HEK293T_min | HEK293T_supp |
|---|---|---|---|---|---|---|
| GO_BP | GO:0000902 | cell morphoge | C7 | 0,0248017 | 0,0181091 | IntAct / MI / Co |
| GO_BP | GO:0006334 | nucleosome a | C4 | 0,0315097 | 0,00023744 | STRING / MI / C |
| GO_BP | GO:0006338 | chromatin rem | C4 | 0,01184382 | 0,00011954 | STRING / MI / C |
| GO_BP | GO:0006399 | tRNA metaboli | C1;C10;C5 | 5,59E-09 | 0,02121524 | IntAct / Pearso |
| GO_BP | GO:0006401 | RNA catabolic | C1 | 0,00096284 | 0,00044824 | IntAct / Pearso |
| GO_BP | GO:0006402 | mRNA catabol | C1 | 0,01889327 | 0,00031984 | IntAct / Pearso |
| GO_BP | GO:0006417 | regulation of t | C1 | 0,00572517 | 0,00191615 | STRING / MI / C |
| GO_BP | GO:0009451 | RNA modificat | C5 | 1,11E-07 | 0,02687438 | STRING / Pears |
| GO_BP | GO:0010608 | post-transcrip | C1 | 0,00378896 | 0,00470797 | STRING / MI / C |
| GO_BP | GO:0017148 | negative regul | C1 | 0,00013792 | 9,94E-05 | STRING / MI / C |
| GO_BP | GO:0022613 | ribonucleopro | C1 | 0,04077497 | 0,0004844 | IntAct / Pearso |
| GO_BP | GO:0030031 | cell projection | C7 | 0,0149185 | 0,00481164 | STRING / Pears |
| GO_BP | GO:0034655 | nucleobase-co | C1 | 0,00396894 | 0,00075416 | IntAct / Pearso |
| GO_BP | GO:0034728 | nucleosome o | C4 | 0,0425175 | 0,00065102 | STRING / MI / C |
| GO_BP | GO:0036257 | multivesicular | C3 | 0,04336867 | 0,00011528 | IntAct / dCor / |
| GO_BP | GO:0036258 | multivesicular | C3 | 0,04336867 | 0,00011528 | IntAct / dCor / |
| GO_BP | GO:0040011 | locomotion | C4 | 0,03785183 | 0,04935687 | IntAct / MI / Co |
| GO_BP | GO:0044782 | cilium organiz | C7 | 0,00831814 | 0,00227749 | STRING / Pears |
| GO_BP | GO:0045947 | negative regul | C1 | 0,02228077 | 2,37E-05 | STRING / MI / C |
| GO_BP | GO:0051205 | protein inserti | C8 | 0,01209555 | 0,00495946 | IntAct / dCor / |
| GO_BP | GO:0051248 | negative regul | C1 | 0,01048729 | 0,00315148 | STRING / MI / C |
| GO_BP | GO:0051668 | localization wi | C7 | 0,00831814 | 0,00314438 | STRING / MI / C |
| GO_BP | GO:0060271 | cilium assemb | C7 | 0,00831814 | 0,00216874 | STRING / Pears |
| GO_BP | GO:0061024 | membrane org | C8 | 2,84E-06 | 0,00480302 | STRING / Pears |
| GO_BP | GO:0061951 | establishment | C7 | 0,00081989 | 0,00027487 | STRING / Pears |
| GO_BP | GO:0072657 | protein localiz | C7;C9 | 0,0149185 | 0,01035487 | STRING / MI / C |
| GO_BP | GO:0072659 | protein localiz | C7;C9 | 0,00831814 | 0,00560094 | STRING / Pears |
| GO_BP | GO:0090150 | establishment | C7 | 0,00081989 | 0,00027487 | STRING / Pears |
| GO_BP | GO:0120031 | plasma memb | C7 | 0,0149185 | 0,00481164 | STRING / Pears |
| GO_BP | GO:0141188 | nucleic acid ca | C1 | 0,00115166 | 0,00044824 | IntAct / Pearso |
| GO_BP | GO:1901699 | cellular respo | C7 | 0,02384626 | 0,03387138 | STRING / MI / C |
| GO_BP | GO:1903313 | positive regula | C1 | 0,0272072 | 0,02046018 | IntAct / Pearso |
| GO_BP | GO:1990778 | protein localiz | C7;C9 | 0,01194398 | 0,00782801 | STRING / Pears |
| Reactome | R-HSA-14451 | Translocation | C7 | 1,74E-06 | 0,00425189 | STRING / Pears |
| Reactome | R-HSA-15922 | Mitochondrial | C8 | 7,69E-08 | 1,27E-10 | STRING / MI / C |
| Reactome | R-HSA-16515 | MTOR signallir | C1 | 0,0224452 | 4,60E-07 | STRING / MI / C |
| Reactome | R-HSA-18522 | Organelle biog | C8 | 9,40E-06 | 3,41E-08 | STRING / MI / C |
| Reactome | R-HSA-19431 | Signaling by Rl | C9 | 0,01684438 | 0,00622057 | IntAct / MI / Co |
| Reactome | R-HSA-19999 | Membrane Tra | C7 | 0,00259407 | 0,00040747 | STRING / Pears |
| Reactome | R-HSA-52509 | Positive epige | C4 | 0,00097004 | 0,01688253 | IntAct / dCor / |
| Reactome | R-HSA-56288 | TP53 Regulate | C1 | 0,04576634 | 2,64E-06 | STRING / MI / C |
| Reactome | R-HSA-56536 | Vesicle-media | C7 | 0,00259407 | 0,00040747 | STRING / Pears |
| Reactome | R-HSA-71291 | Metabolism of | C6 | 0,00018385 | 0,00866237 | STRING / Pears |
| Reactome | R-HSA-72766 | Translation | C10 | 3,00E-06 | 0,0403164 | STRING / Pears |
| Reactome | R-HSA-88761 | RAB GEFs excl | C7 | 0,02171168 | 1,45E-08 | STRING / Pears |
| Reactome | R-HSA-88763 | Listeria mono | C9 | 0,03143018 | 0,00630175 | IntAct / dCor / |

| | | | | | | |
|---|---|---|---|---|---|---|
| Reactome | R-HSA-89496[illegible] | Cristae format | C8 | 1,07E-09 | 1,59E-13 | STRING / MI / [illegible] |
| Reactome | R-HSA-90071[illegible] | Rab regulation | C7 | 0,03365129 | 3,78E-07 | STRING / Pears |
| Reactome | R-HSA-90129[illegible] | RHO GTPase c | C9 | 0,03626108 | 0,02624883 | IntAct / MI / Co |
| Reactome | R-HSA-90134[illegible] | RHOH GTPase | C6 | 0,04113794 | 0,04089001 | STRING / MI / [illegible] |
| Reactome | R-HSA-97165[illegible] | Signaling by RI | C9 | 0,01684438 | 0,00622057 | IntAct / MI / Co |
| Reactome | R-HSA-98566[illegible] | MITF-M-depen | C1 | 0,04449726 | 0,00140654 | STRING / MI / [illegible] |

mponent_001 / adjusted P=0,0346646387315564; STRING / Pearson / Component_003 / adjusted P=0,
Component_001 / adjusted P=0,000237435819779044; STRING / dCor / Component_001 / adjusted P=0
Component_001 / adjusted P=0,000119544387099909; STRING / dCor / Component_001 / adjusted P=0

son / Component_002 / adjusted P=0,0342189321672837; STRING / dCor / Component_003 / adjusted

n / Component_002 / adjusted P=0,0240104319448328; STRING / Pearson / Component_002 / adjusted
son / Component_003 / adjusted P=0,00481163702234781; STRING / dCor / Component_004 / adjusted

Component_001 / adjusted P=0,000651019359326533; STRING / dCor / Component_001 / adjusted P=0
Component_002 / adjusted P=0,0169472227856653; STRING / ConstMI / Component_002 / adjusted P=
Component_002 / adjusted P=0,0169472227856653; STRING / ConstMI / Component_002 / adjusted P=

son / Component_003 / adjusted P=0,00227749470382785; STRING / dCor / Component_004 / adjusted

Component_003 / adjusted P=0,0338713796389636; STRING / Pearson / Component_003 / adjusted P=
son / Component_003 / adjusted P=0,00216873989344215; STRING / dCor / Component_004 / adjusted
son / Component_003 / adjusted P=0,0142341958112707; STRING / dCor / Component_004 / adjusted
son / Component_003 / adjusted P=0,000274873238179593; STRING / dCor / Component_004 / adjuste
Component_003 / adjusted P=0,0141626773805083; STRING / Pearson / Component_003 / adjusted P=
son / Component_003 / adjusted P=0,00560094346108819; STRING / dCor / Component_004 / adjusted
son / Component_003 / adjusted P=0,000274873238179593; STRING / dCor / Component_004 / adjuste
son / Component_003 / adjusted P=0,00481163702234781; STRING / dCor / Component_004 / adjusted

son / Component_003 / adjusted P=0,00782800989525468; STRING / dCor / Component_004 / adjusted
son / Component_003 / adjusted P=0,00425189428070084; STRING / dCor / Component_004 / adjusted
Component_002 / adjusted P=3,59310844548796e-08; STRING / Pearson / Component_001 / adjusted P

Component_002 / adjusted P=8,80266427374784e-06; STRING / Pearson / Component_001 / adjusted P

son / Component_003 / adjusted P=0,00257322995693969; STRING / dCor / Component_004 / adjusted

son / Component_003 / adjusted P=0,00257322995693969; STRING / dCor / Component_004 / adjusted

son / Component_003 / adjusted P=3,89883326809043e-06; STRING / dCor / Component_004 / adjuste

Component_002 / adjusted P=4,63979066839063e-11; STRING / Pearson / Component_001 / adjusted P
son / Component_003 / adjusted P=3,11906661447235e-05; STRING / dCor / Component_004 / adjuste

Component_002 / adjusted P=0,0014065400266509; STRING / Pearson / Component_001 / adjusted P=

018109095483321; STRING / dCor / Component_004 / adjusted P=0,033192157224792

P=0,00928467613699087; STRING / dCor / Component_003 / adjusted P=0,000484395551455001

0,00314438226928242; STRING / dCor / Component_004 / adjusted P=0,00613315278115081; STRIN

P=0,0252712356566047; STRING / ConstMI / Component_002 / adjusted P=0,00480301824913832

0,0103548671699804; STRING / dCor / Component_004 / adjusted P=0,0162209346522737

=3,59310844548796e-08; STRING / dCor / Component_002 / adjusted P=1,27354455716533e-10; STR

=8,80266427374784e-06; STRING / Pearson / Component_003 / adjusted P=0,0179334684466554; ST [illegible]=4,63979066839063e-11; STRING / dCor / Component_002 / adjusted P=1,58961251154399e-13; STR[illegible]

0,0014065400266509; STRING / ConstMI / Component_001 / adjusted P=0,00156896207679989; STRI[illegible]

'RING / dCor / Component_002 / adjusted P=3,41066306709372e-08; STRING / dCor / Component_004

/ adjusted P=0,0346016763211678; STRING / ConstMI / Component_001 / adjusted P=0,00017477054[illegible]

3986391
